# Anti-seed PNAs targeting multiple oncomiRs for brain tumor therapy

**DOI:** 10.1101/2022.01.31.478549

**Authors:** Yazhe Wang, Shipra Malik, Hee-Won Suh, Yong Xiao, Yanxiang Deng, Rong Fan, Anita Huttner, Ranjit S Bindra, W Mark Saltzman, Raman Bahal

## Abstract

Glioblastoma (GBM) is one of the most lethal malignancies in the United States with poor survival and high recurrence rates, suggesting the need for approaches targeting the most important molecular drivers of tumor growth. Here, we aimed to simultaneously target oncomiRs 10b and 21, which have been reported to drive the aggressive growth and invasiveness of GBM. We designed **s**hort (8-mer bases) gamma-(**γ**)-modified **p**eptide **n**ucleic **a**cids (sγPNAs), which target the seed region of oncomiRs 10b and 21 with high affinity. We entrapped these anti-miR sγPNAs in nanoparticles (NPs) formed from a block copolymer of poly(lactic acid) and hyperbranched polyglycerol (PLA-HPG); the NPs were also functionalized with aldehydes to produce bioadhesive NPs. We have previously shown that these bioadhesive NPs (BNPs) produce superior transfection efficiency, with a tropism for tumor cells. The sγPNA BNPs showed superior anti-miR efficacy in comparison to the regular full length PNA BNPs *in vitro*. When combined with temozolomide, sγPNA BNPs administered via convention-enhanced delivery (CED) inhibited the growth of intracranial tumors and significantly improved the survival of animals (>120 days). RNA sequencing analysis revealed the role of vascular endothelial growth factor A (VEGFA) and integrin beta 8 (ITGB8), direct targets of both miR-10b and miR-21, in mediating the tumor growth. Hence, we established that BNPs loaded with anti-seed sγPNAs targeting multiple oncomiRs is a promising approach to improve the treatment of GBM, with a potential to personalize treatment based on tumor specific oncomiRs.

**Summary:** *Targeting oncomiRs 21 and 10b to improve GBM survival:* Glioblastoma (GBM) is an aggressive malignant disorder with high recurrence rates and poor survival. Here, we aimed to simultaneously inhibit two aberrant oncomiRs—miR 21 and miR 10b—which have been previously associated with GBM invasiveness and progression. We synthesized short, gamma-modified peptide nucleic acids (sγPNA) targeted to the miR seed regions and loaded the sγPNAs into bioadhesive nanoparticles (BNPs). When the sγPNA-BNPs were added to cultured tumor cells, we observed significant reduction of target oncomiRs and increase of apoptosis *in vitro*. When delivered *in vivo* by convection-enhanced delivery, sγPNA BNPs dramatically increased the survival in two orthotopic (intracranial) mouse models of GBM. Moreover, the combination of sγPNA BNPs with temozolomide (TMZ) increased the survival of mice with GBM beyond the planned endpoint (120 days) with significant improvements in histopathology. The proposed strategy of sγPNA BNP with TMZ provides an alternative, promising approach for treatment of GBM.

## INTRODUCTION

Glioblastoma (GBM) is the most aggressive form of brain tumor, affecting 70-75% of adults with high mortality and median survival of only 14-17 months (*1*). GBM has an alarmingly high incidence rate of 4.32 per 100,000 in the United States (*2*) and a one year survival rate of only 1.4% in patients over 75 years of age (*3*). The primary therapeutic approach for GBM is surgical resection followed by radiotherapy and chemotherapy. Temozolomide (TMZ) improves the two-year survival rate from 10.4% to 26.5% in combination with radiotherapy, compared to radiotherapy alone (*4*). Hence, there is a need for improved therapeutic approaches for targeting molecular genetic mediators of GBM.

MicroRNAs (miRNAs or miRs) are ∼20-25 nucleotide non-coding RNAs that regulate gene expression at the post-transcriptional level (*5*). The dysregulation of miRs, either upregulation (where they are known as oncomiRs) (*6*) or downregulation, plays an important role in several malignancies (*7–10*). Previous studies have reported aberrant miR expression levels in GBM patients; some of these are associated with poor prognosis and low overall survival (*11*). In particular, miR-10b (*12–15*) and miR-21 (*16–20*) appear to be the most highly upregulated oncomiRs contributing to GBM. miR-10b enhances GBM growth by negatively regulating Bim (BCL2 interacting mediator of death), TFAP2C (transcription factor AP-2γ), CDKN2A/16 (tumor suppressor), and p21 (cell cycle inhibitor) expression (*13*). Inhibiting miR-10b reduces the growth of intracranial GBM tumors in animal models (*14, 21*). These promising results have prompted the development of an investigational antisense oligonucleotide (RGLS5799, Regulus Therapeutics) targeting miR-10b. Similarly, upregulated miR-21 levels increase GBM invasiveness by inhibiting matrix metalloproteinase (MMP) (*17*), induce proliferation via negative regulation of insulin-like growth factor binding protein-3 (IGFBP3) (*20*) or phosphatase and tensin homolog (PTEN), and promote tumor stemness via SOX-2, a transcription factor (*22, 23*). Hence, knocking down miR-21 reduces GBM progression and invasion (*24–26*) in addition to preventing chemoresistance of GBM cells to TMZ (*27, 28*) and taxol (*29*). Current therapeutic strategies, which are focused on targeting a single oncomiR, have shown limited efficacy against GBM.

In this study, we aimed at targeting miR-10b and miR-21 simultaneously to extend survival and to enhance the chemosensitization of GBM towards TMZ. PNAs are synthetic nucleic acid analogues where the phosphodiester backbone is substituted with neutral N-(2-aminoethyl) glycine units (*30*). PNAs bind to target miRs via complementary Watson and Crick base pairing, but they are enzymatically stable (*31, 32*). Classical PNAs targeting full length oncomiRs have been explored for cancer therapy (*33–37*), but the functional activity of miRs is governed by the “seed region” centered on nucleotides 2 to 7 on the 5’ end (*38*). Hence, antimiR efficacy can be achieved by targeting only the seed region of oncomiRs (*39*). Here, we designed serine-gamma PNAs (γPNAs) complementary to the seed region of oncomiR-21 and oncomiR-10b to achieve improved antimiR activity. Because γPNAs are pre-organized into a helical conformation due to the presence of chirality at γ position, they have superior physicochemical features, binding affinity, and specificity compared to classical PNAs (*40–42*), making it possible to target short sequences with high affinity. Short anti-seed γPNAs (sγPNAs) possess numerous appealing features: synthesis is straightforward, quality control analysis is simplified over longer sequences, and they are well suited for conjugation with fluorophores or other entities to enable imaging. Most importantly, anti-seed sγPNAs are clinically more translatable and possess comparable *in vivo* efficacy to conventional full-length anti-miRs with minimal toxicity (*39, 43–45*).

Over the past years, we have developed an approach for delivery of agents directly to brain tumors, using convection-enhanced delivery (CED) to introduce polymer NPs, that are loaded with active agents, directly into the brain (*46*). NPs composed of a block copolymer of poly (lactic acid) and hyperbranched polyglycerol (PLA-HPG), which can be surface functionalized to introduce aldehyde groups (*47*), creating PLA-HPG-CHO, have several advantages for delivery of PNA anti-miRs. NPs formed from PLA-HPG-CHO are bioadhesive nanoparticles, or BNPs (*48–51*). We have shown that—compared with several other NPs of similar composition—these BNPs lead to the highest levels of uptake into intracranial tumor cells after CED (*52*) and they can be loaded with PNAs, which are slowly released after CED in the brain (*53*). In this work, we loaded the PLA-HPG-CHO BNPs with two sγPNAs—one binding to miR-10b and another binding to miR-21—to test the hypothesis that simultaneous delivery of two anti-miRs in glioma cells can regulate unique molecular pathways leading to improved survival.

## RESULTS

### Design and synthesis of PNA oligomers

Regular and serine-γPNA oligomers targeting oncomiR-21 and oncomiR-10b were synthesized on a solid support using standard Boc chemistry protocols (Fig. 1) (*54*). sγPNA-21 and sγPNA-10b are sγPNAs—with a serine modification at the γ-position—designed to bind the seed region of oncomiRs 21 and 10b, respectively. Further, sγPNA-21 and sγPNA-10b were extended with three arginine residues on the N-terminus to increase the binding to their respective targets via electrostatic interaction between the cationic domain of γPNAs and the negatively charged backbone in the flanking region of miRNA (*55, 56*). Scrambled versions of these PNAs—Scr-sγPNA-21 and Scr-sγPNA-10b—were used as controls. To compare the efficacy of short cationic γPNAs with full length regular PNAs, we also synthesized PNA-21 and PNA-10b, which were designed to target full length oncomiRs 21 and 10b, respectively (Fig. 1). TAMRA dye (5-carboxytetramethylrhodamine) was conjugated to the N-terminus of the PNAs for visualization of cellular uptake and biodistribution. Reverse phase high performance liquid chromatography (RP-HPLC) and mass spectrometry analysis confirmed the high quality of the synthesized PNA preparations (Fig. S1 and Table S1).

**Fig. 1.**
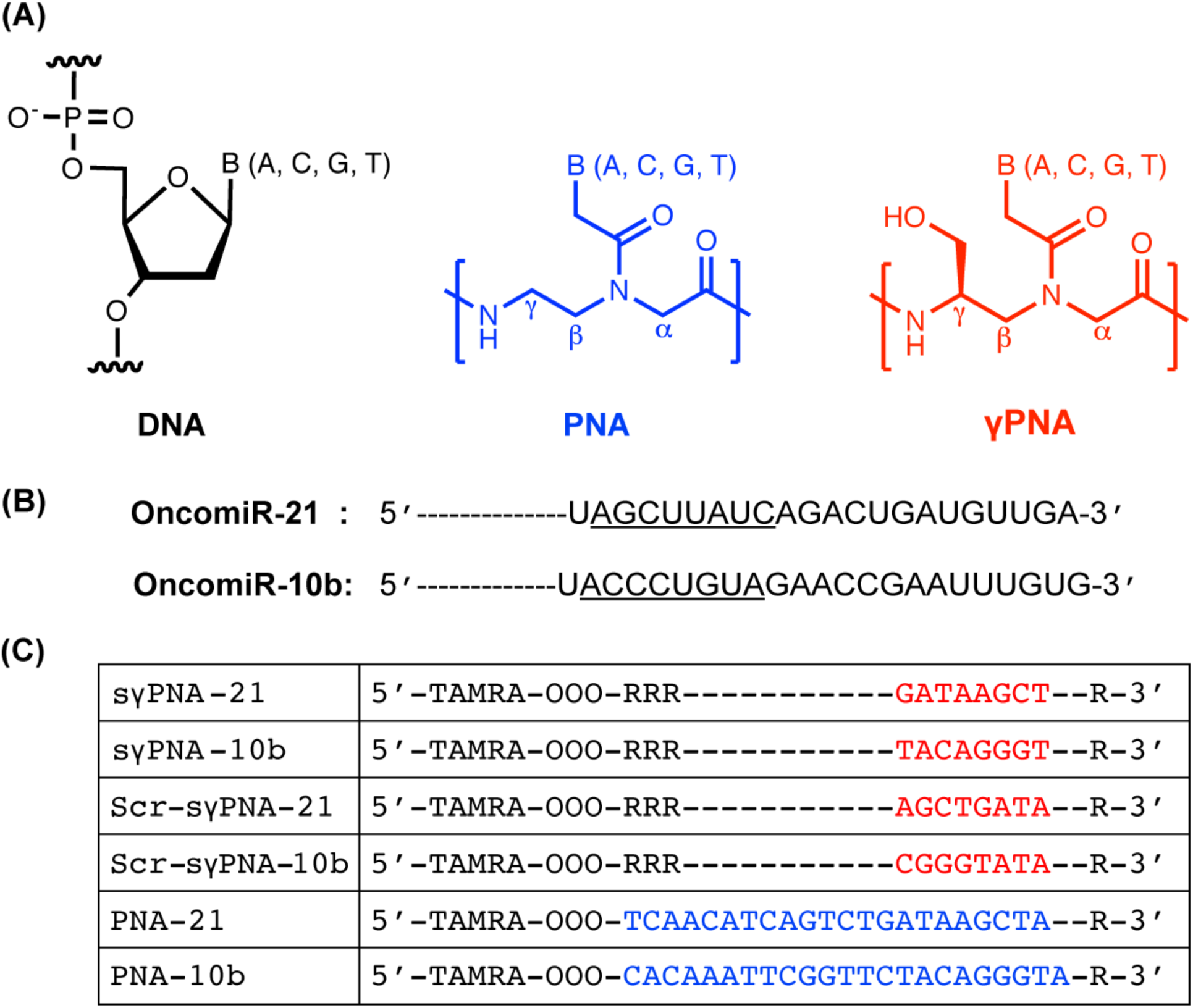
The sequence of Peptide Nucleic Acids (PNAs) used in the study. **(A)** Chemical structures of DNA, PNA and serine-γ-PNA. **(B)** The sequence of oncomiR-21 and oncomiR-10b, where the underlined segment is the seed region. **(C)** PNAs used in this study. sγPNA-21 and sγPNA-10b are serine-γPNAs designed to bind the seed region of oncomiR-21 and oncomiR-10b, respectively. Three arginine (RRR) residues are appended to the N-terminus and one arginine (R) on the C-terminus. Scr-sγPNA-21 and Scr-sγPNA-10b are scrambled versions of sγPNA-21 and sγPNA-10b, respectively. PNA-21 and PNA-10b are regular PNAs designed to bind full length oncomiR-21 and oncomiR-10b, respectively. PNAs were conjugated with 5-carboxytetramethylrhodamine (TAMRA), a fluorescent dye for imaging. OOO represents 8-amino-2,6,10-trioxaoctanoic acid residues (Mini-PEG). There are present to form a flexible linker connecting the TAMRA and Watson Crick binding regions of the PNAs.

### sγPNAs bind to the target oncomiRs with high affinity and specificity

Binding affinity was measured by incubating the PNAs with the target oncomiRs. sγPNA-21 and sγPNA-10b were incubated with miR-21 and miR-10b, respectively, in simulated physiological conditions at two different ratios of 2:1 and 4:1 (PNA:miR) for 16 h. Both sγPNA-21 and sγPNA-10b showed significant binding to their respective targets even at the lower 2:1 ratio (Fig. S2A), as evidenced by the faint band corresponding to the unbound miRs and the prominent retarded band of the PNA-miR heteroduplexes. The band for target miR-21 and miR-10b completely disappeared on incubation with their respective PNAs at 4:1 ratio (Fig. S2B). We also studied the binding of sγPNA-21 and sγPNA-10b in the presence of both miR-10b and miR-21. When incubated with the target miR-21 at 1:1 ratio, sγPNA-21 showed a retarded band (Fig. S3, lane 3). As expected, on incubation of sγPNA-21 with miR 10b, we did not observe any shift in the band (Fig. S3, lane 6). Incubation of sγPNA-21 with both the miRs resulted in only one retarded band (Fig. S3, lane 7) similar to the sγPNA-21-miR-21 heteroduplex, indicating the specificity of sγPNA-21 towards the target miR-21. Similarly, sγPNA-10b showed a shifted band only after incubation with the target miR-10b (Fig. S3, lane 4) and no retarded band was visible in presence of miR-21 (Fig. S3, lane 5). Further, on incubation with both the miRs, sγPNA-10b showed one retarded band similar to the sγPNA-10b-miR-10b heteroduplex. Hence, sγPNA-21 and sγPNA-10b showed high binding affinity as well as specificity towards miR-21 and miR-10b, respectively.

### Preparation and characterization of nanoparticles

PNAs were encapsulated in PLA-HPG to produce non-adhesive NPs (NNPs), using a modified single-emulsion method as previously reported (*47, 53*). BNPs were prepared by brief exposure of these NNPs to sodium periodate, converting the vicinal diols of HPG to aldehydes, creating HPG-CHO; successful conversion to the bioadhesive state was confirmed by measuring BNP adhesion to poly(L-lysine) coated glass (Fig. S4). For the simultaneous knockdown of target oncomiRs 10b and 21, different batches of NPs formulated with respective PNAs were physically mixed at a fixed PNA molar ratio (1:1). Single sγPNA loaded NPs and combinations of NP formulations (sγPNA/NNP: sγPNA/PLA-HPG and sγPNA/BNP: γPNA/PLA-HPG-CHO) were extensively characterized (Table 1). sγPNA/NNP and sγPNA/BNP had similar average hydrodynamic sizes, ranging from 150 to 165 nm, as measured by dynamic light scattering (DLS). All the NP formulations exhibited negative surface charge in water with ζ potential between -20 mV to -30 mV. The average PNA loading was around 1.5 nmol per mg of NPs suggesting superior encapsulation efficiency. TEM images showed uniform and spherical morphology of sγPNA/BNP and sγPNA/NNP (Fig. 2A). The majority of visible NPs were around 100 nm in diameter, which confirmed the desired particle size. Both sγPNA/NNP and sγPNA/BNP were stable with no measurable aggregation during 3 days of incubation in artificial CSF (aCSF) (Fig. 2B). *In vitro* sγPNA release from both NNP and BNP during continuous incubation in PBS at 37°C was similar (Fig. 2C), with slightly slower release from BNPs.

**Fig. 2.**
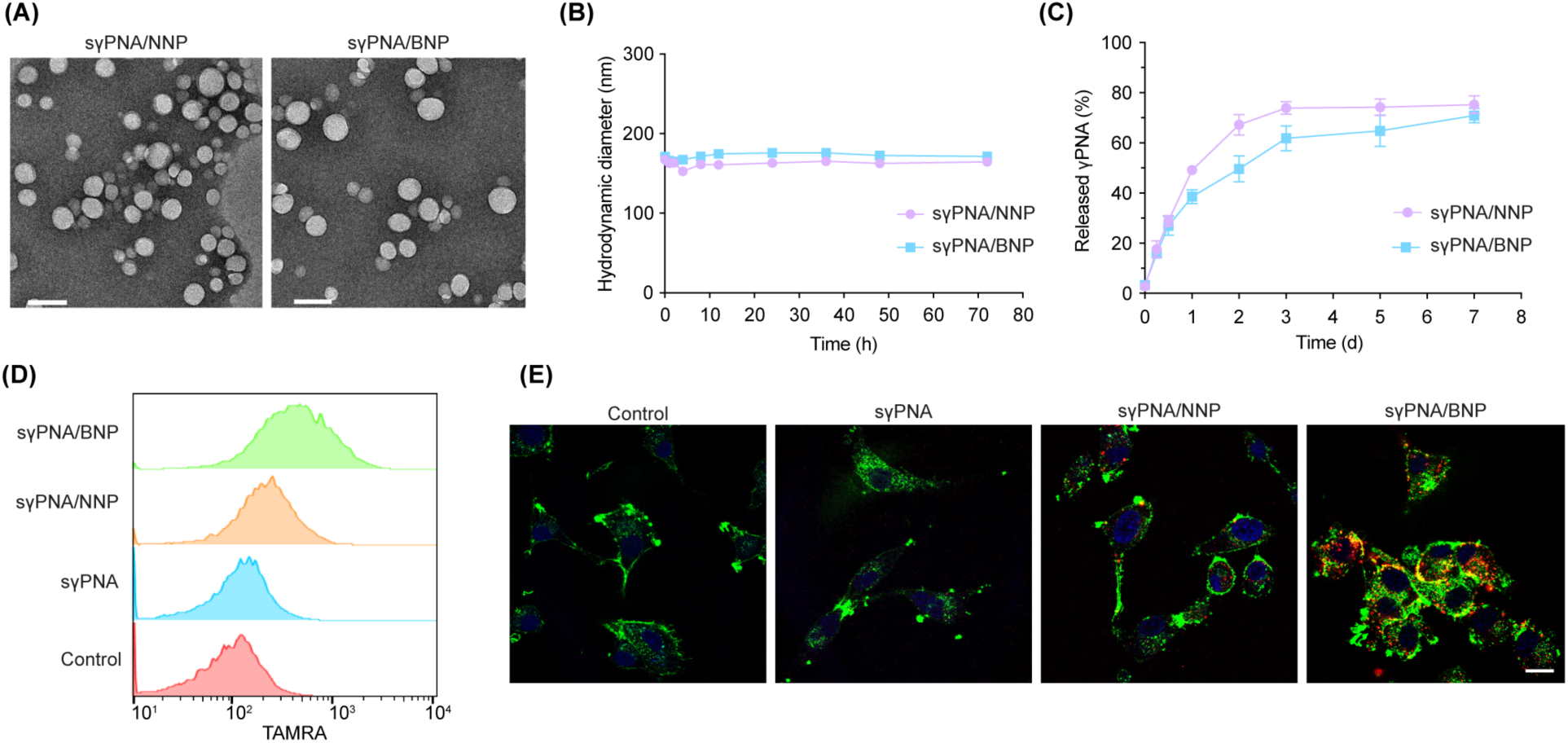
Characterization of sγPNA loaded nanoparticles. **(A)** TEM images of sγPNA/NNP and sγPNA/BNP. Scale bar, 100 nm. **(B)** Size stability of sγPNA/NNP and sγPNA/BNP in aCSF. **(C)** The amount of sγPNA (sγPNA-21 and sγPNA-10b) released from NPs over time during incubation in buffered saline was determined and quantified as a percentage of amount loaded. The data are shown as mean ± SD (n = 3). **(D)** Cellular uptake analyzed by flow cytometry. Top to bottom: U87 cells treated with γPNA/BNP, γPNA/NNP, free γPNA (physical mixture of free γPNA1 and γPNA2) and untreated cell. **(E)** Confocal microscopic images of cells treated by free γPNA or γPNA loaded NPs. PNAs were conjugated with TAMRA (red), F-actin was labelled with phalloidin (green) and nucleus was stained with Hoechst (blue). Scale bar, 20 µm. sγPNA/BNP are physical mixture of sγPNA-21 BNP and sγPNA-10b BNP. sγPNA/NNP are physical mixture of sγPNA-21 NNP and sγPNA-10b NNP. NNP indicates PLA-HPG nanoparticles and BNP indicates PLA-HPG-CHO nanoparticles.

**Table 1.**
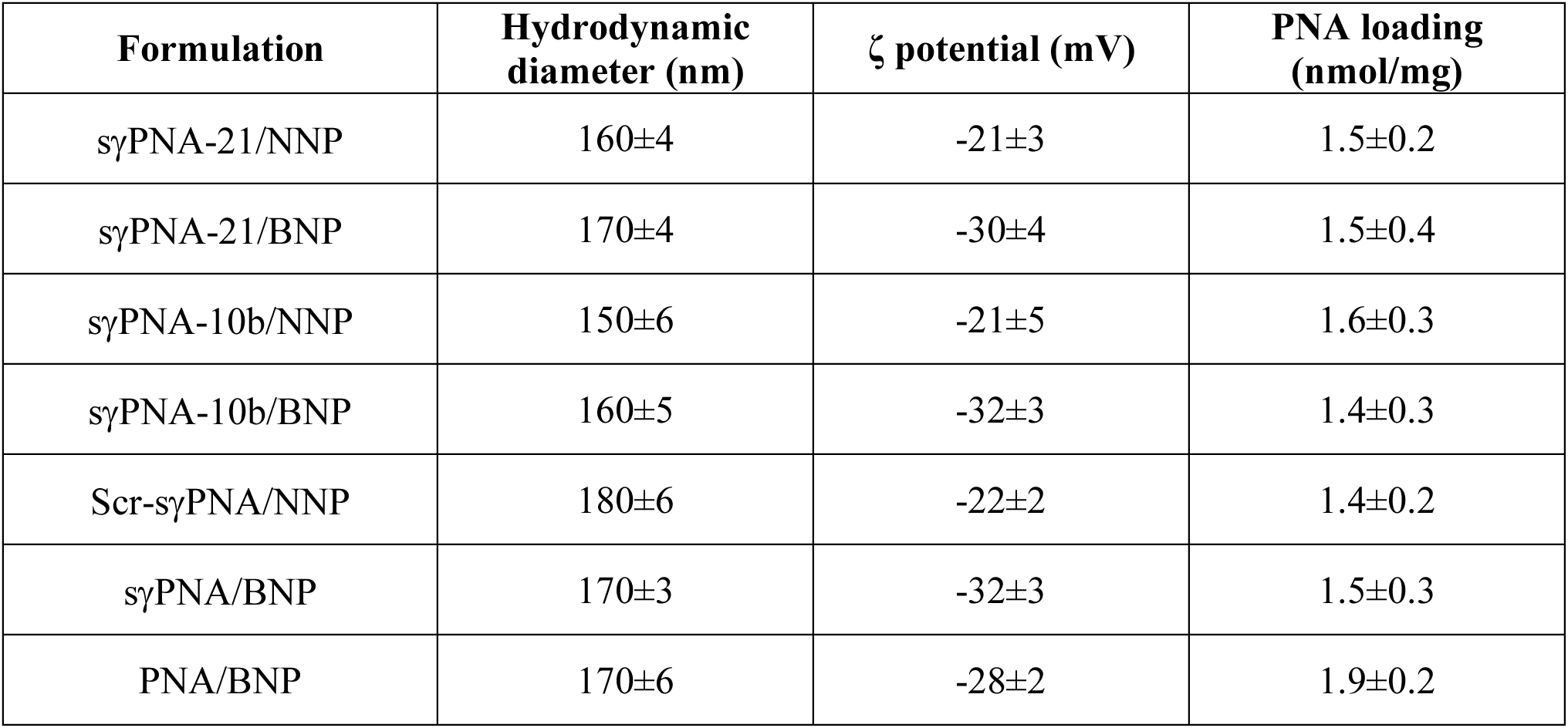
Hydrodynamic diameter, ζ potential and PNA loading in various nanoparticles. The data are shown as mean ± SD (n = 3). Scr-sγPNA/NNP was the physical mixture of Scr-sγPNA-21 loaded NNP and Scr-sγPNA-10b loaded NNP, sγPNA/BNP was the mixture of sγPNA-21 loaded BNP and sγPNA-10b loaded BNP, PNA/BNP was the mixture of PNA-21 loaded BNP and PNA-10b loaded BNP. NNP indicates PLA-HPG nanoparticles and BNP indicates PLA-HPG-CHO nanoparticles.

### sγPNA loaded BNPs show superior cellular uptake in glioma cells

TAMRA fluorescence from sγPNAs was used to quantify the uptake of NPs in glioma cells. Flow cytometry analysis revealed that the uptake of BNP was significantly higher than that of NNP in U87 cells (Fig. 2D and S5A). Free sγPNAs showed no significant uptake after 24 h incubation. For microscopic observation, cells treated with either NNP or BNP showed stronger TAMRA fluorescence than cells treated with free sγPNA. Confocal microscopy (Fig. 2E) indicated that treatment with BNP led to higher cellular uptake than NNP; BNP also produced wider distribution throughout the cells. Similar results were obtained in a different human glioma cell line, LN-229 (Fig. S5B). These findings confirm the preferential cellular uptake of sγPNA/BNP into glioma cells, which is consistent with our previous studies (*26, 52*).

### sγPNA loaded BNP inhibit expression of miR-10b and miR-21

To confirm the therapeutic efficacy of designed anti-miR sγPNAs, different types of PNA-loaded NP formulations or free sγPNAs were incubated with U87 cells. Cellular levels of miR-21 and miR-10b were significantly reduced 72 h after treatment with NPs (Fig. 3A). A significant decrease of miR-10b expression was achieved following incubation with sγPNA/NNP and sγPNA/BNP when compared to the control treatment (Fig. 3A); BNP also produced a significant decrease in miR-21 expression after treatment, while NNP exhibited a slight but not statistically significant effect. Scrambled versions of sγPNA NP formulations and free sγPNAs had no effect on the miR-10b and miR-21 expression (Fig. 3A). Of note, sγPNA/BNP showed greater suppression of both oncomiRs than full length PNA loaded NPs (PNA/BNP), which we believe reflects the higher binding affinity of sγPNAs (Fig. 3B). We found that PTEN, one of the most frequently mutated tumor suppressor genes in human cancer (*57*) and a predicted target of both miR-21 (*58*) and miR-10b (*14*), was upregulated, by 4-5 fold after treatment with sγPNA loaded NPs (Fig. S6A). Moreover, the specific inhibition of miR-21 mediated by sγPNA-21 loaded NPs did not affect the expression of miR-10b and, likewise, sγPNA-10b loaded NPs did not affect the expression of miR-21 (Fig. S6B). These results demonstrated the specific and effective knockdown of target oncomiRs provided by sγPNA loaded NPs, with BNPs providing the strongest biological effect.

**Fig. 3.**
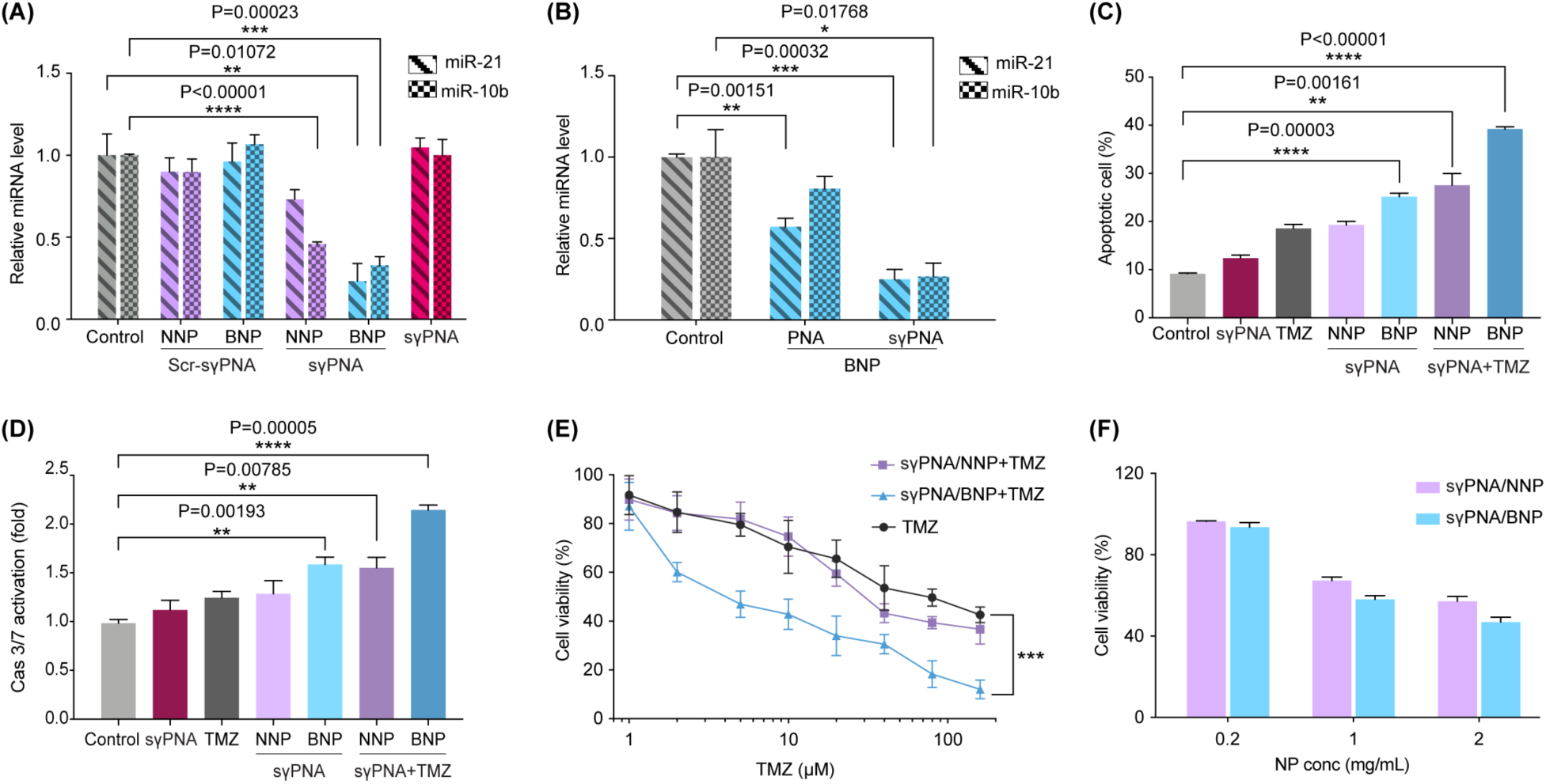
sγPNA mediated simultaneous knockdown of miR-21 and miR-10b induces apoptosis in glioma (U87) cells. **(A)** Expression analysis of miR-10b and miR-21 levels in U87 cells after treatment with Scr-sγPNA/NNP, Scr-sγPNA/BNP, sγPNA/NNP, sγPNA/BNP, and free sγPNA. Scr-sγPNA/NNP and Scr-sγPNA/BNP indicates the physical mixture of Scr-sγPNA-21 and Scr-sγPNA-10b loaded NPs (NNP or BNP). sγPNA/NNP and sγPNA/BNP indicates the physical mixture of sγPNA-21 and sγPNA-10b loaded NPs (NNP or BNP). Data are expressed as mean ± SD (n = 3). **(B)** Relative miRNA level in U87 cells treated with different formulations. PNA/BNP was the physical mixture of PNA-21 loaded BNP and PNA-10b loaded BNP. **(C)** Percentage of early apoptotic cells after different treatments with sγPNA/NNP and BNP with or without temozolomide (TMZ). Data are expressed as mean ± SD (n = 3). **(D)** Caspase 3 and caspase 7 activities in cells treated by combinations of NPs and TMZ for 72 h. Data are expressed as mean ± SD (n = 3). **(E)** Viability of cells treated with increasing doses of TMZ, with or without sγPNA NPs (NNP or BNP) treatment. **(F)** Viability of cells treated with different doses of NPs for 72 h.

### sγPNA loaded BNPs induce enhanced apoptosis of U87 cells in combination with temozolomide

Apoptosis was measured by annexin V and PI staining in cells after treatment with sγPNA loaded NPs. Apoptosis was enhanced by sγPNA/BNP treatment (compared to sγPNA/NNP-treated or control cells), and further enhanced after combined treatment with TMZ (Fig. 3C, Fig. S7). Co-treatment with TMZ and sγPNA/BNP led to higher apoptotic activity than full length PNA BNP (Fig. S8A). Caspase 3 and caspase 7 activities—key indicators of apoptosis—were elevated 2-fold after TMZ and sγPNA/BNP co-treatment (Fig. 3D, S8B).

### Tumor cell death by combination treatment with TMZ

Recent studies show that miRNAs play a role in TMZ resistance, and that regulation of miRNA can enhance TMZ-induced cell death (*59*). We found treatment of U87 cells with sγPNA-loaded NPs sensitized the cells to TMZ (Fig. 3E). Both sγPNA/NNP and sγPNA/BNP showed dose-dependent cytotoxicity against U87 cells (Fig. 3F, S8C). sγPNA/BNP displayed higher cell killing activity and resulted in up to 54% of cell death after treatment for 72 h, while sγPNA/NNP induced 43% cell death at the same condition. The oncogenic miRNA-targeting combination treatment did not produce significant toxicity on human astrocytes (Fig. S8D).

### Simultaneous inhibition of miR-10b and miR-21 decreases VEGFA and ITGB8 levels

To identify the cellular pathways associated with inhibition of tumor growth, we performed RNA sequencing in sγPNA/BNP treated U87 cells (Fig. S9). Upregulated and downregulated genes were identified in treated U87 cells in comparison to the control, using fold change values of greater than 1.5 with significance (padj) value <0.05 (Fig. S10). The hierarchical clustering of log transformed fold change values of differentially and significantly expressed genes (Fig. 4A) revealed a substantial number of dysregulated genes after knockdown of oncomiRs 21 and 10b in comparison to knockdown of each individual oncomiR (Fig. S11 and S12). Gene ontology analysis in U87 cells after the knockdown of oncomiR 21 indicated enrichment of differentially expressed genes (DEGs) associated with PI3-Akt and focal adhesion pathway (Fig. S13) while oncomiR 10b knockdown resulted in enrichment of only PI3-Akt pathway (Fig. S14). But we observed the highest enrichment of DEGs in three major pathways; PI3-Akt, HIF-1, and focal adhesion, involved in angiogenesis, proliferation, survival, and metastasis of tumor cells (Fig. 4B) after knockdown of oncomiRs 10b and 21 in U87 glioma cells. Next, we isolated the DEGs in each of the enriched pathways, including PI3-Akt, HIF, and focal adhesion after knockdown of oncomiR 21 and 10b (Fig. 4C, 4D, and 4E). Further, to isolate the genes associated with GBM pathology and directly regulated targets via miR-10b and miR-21, we intersected the observed DEGs with their predicted targets, i.e., known targets of both miR-10b and miR-21. We obtained 21 direct targets of both miR-21 and miR-10b from the downregulated DEGs identified in the sγPNA/BNP treated group (Fig. 5A). Further intersection of direct downstream targets of miR-10b and miR-21 with the enriched pathways (Fig. 5B, 5C, 5D) revealed the role of vascular endothelial growth factor A (VEGFA) in all three pathways, i.e., PI3-Akt, HIF-1, and focal adhesion. In addition, integrin beta protein 8 (ITGB8) was also identified as a significant regulator of PI3-Akt and focal adhesion pathway. We further validated these findings by gene expression analysis of VEGFA and ITGB8 levels where we noted significant downregulation of both genes in sγPNA/BNP treated U87 cells (Fig. 5E).

**Fig. 4.**
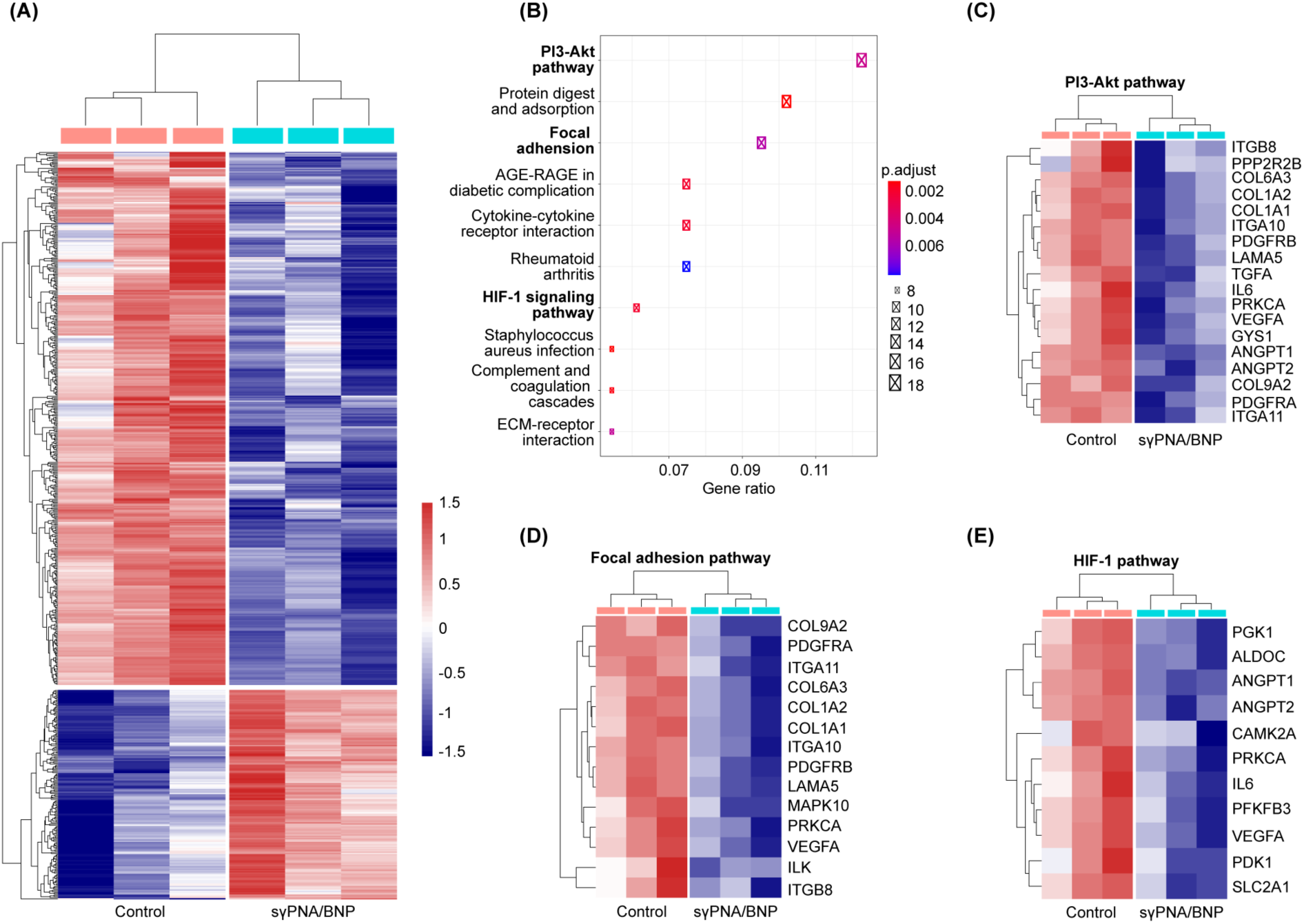
RNA sequencing analysis of sγPNA/BNP treated U87 cells. **(A)** Hierarchal clustering analysis of differentially expressed genes (DEGs) in U87 cells treated with sγPNA/BNP in comparison to control (untreated U87 cells). **(B)** Gene ontology analysis of differentially expressed downregulated genes. **(C)** Heatmap of DEGs associated with PI3-Akt pathway. **(D)** Heatmap of DEGs associated with the focal adhesion pathway. **(E)** Heatmap of DEGs associated with the HIF-1 pathway. sγPNA/BNP indicates the physical mixture of sγPNA-21 and sγPNA-10b loaded BNP. BNP indicates PLA-HPG-CHO nanoparticles.

**Fig. 5.**
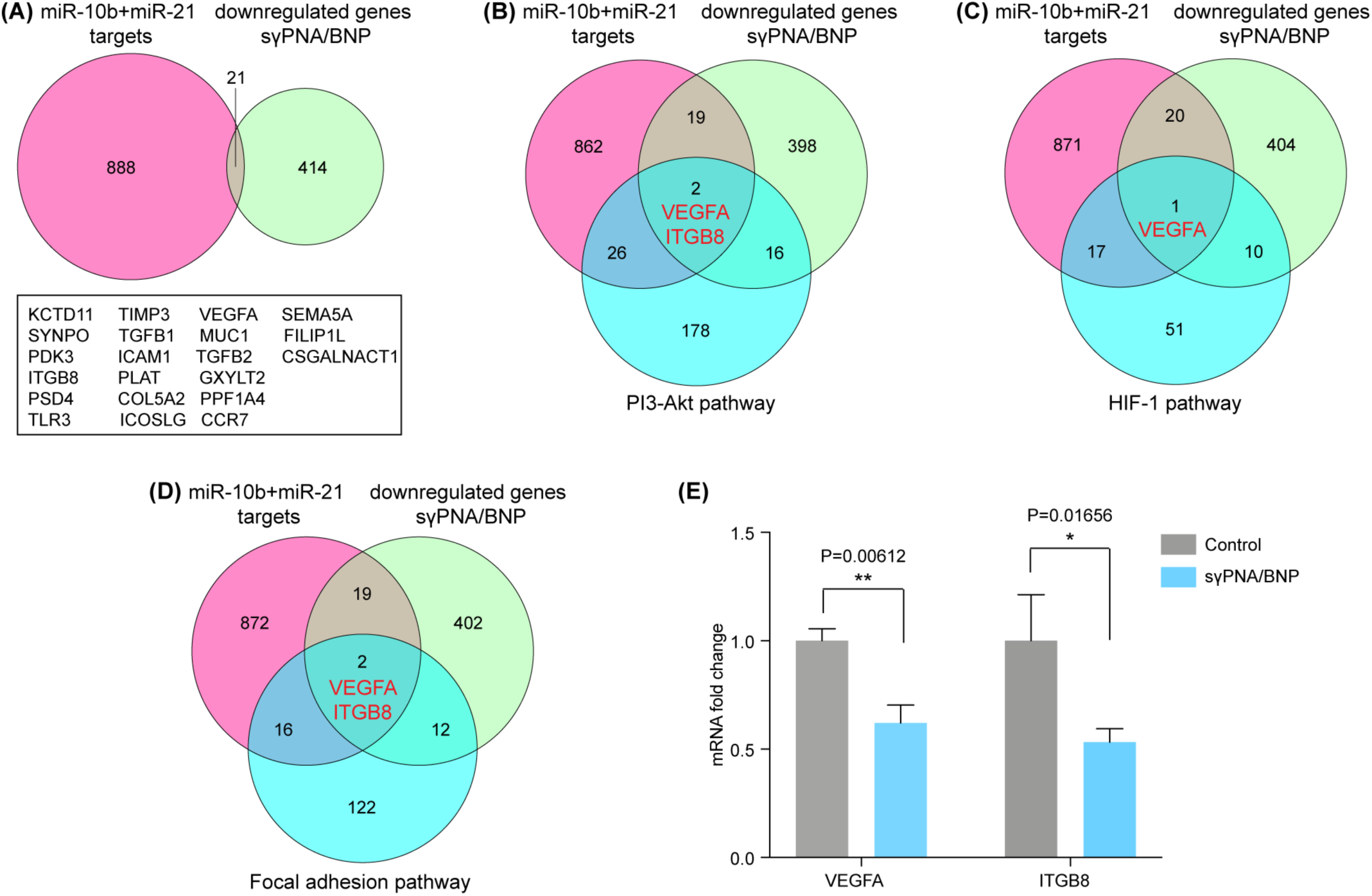
Identifying downstream targets after simultaneous knockdown of miR-21 and miR-10b. **(A)** Venn diagram representing the correlation between predicted miR-10b and miR-21 targets with downregulated DEGs in sγPNA/BNP treated U87 cells. Venn diagram representing the correlation between predicted miR-10b and miR-21 targets with downregulated DEGs in sγPNA/BNP treated U87 cells and genes associated with **(B)** the PI3-Akt pathway, **(C)** the HIF-1 signaling pathway, and **(D)** the focal adhesion pathway. **(E)** Gene expression analysis of VEGFA and ITGB8 in sγPNA/BNP treated U87 cells. sγPNA/BNP indicates the physical mixture of sγPNA-21 and sγPNA-10b loaded BNP. BNP indicates PLA-HPG-CHO nanoparticles.

Finally, to connect our findings with GBM in humans, we correlated survival of GBM patients with overexpression of miR-10b and miR-21. We found poor survival of patients with higher miR-21 levels, but no impact of miR-10b levels on the survival of GBM patients (Fig. S15). However, when patients with higher expression of both miR-10b and miR-21 were examined, the survival probability reduced to less than half when compared with low levels of miR-10b and miR-21 (Fig. S16). These results indicate that upregulation of both miR-10b and miR-21 contribute towards aggressive growth and poor survival in GBM. Hence, the proposed strategy of targeting multiple oncomiRs can pave way for novel personalized therapies for GBM.

### Tumor retention of sγPNA loaded BNP after CED

To study tumor retention and biodistribution of sγPNA loaded in BNP after CED, we infused TAMRA-labeled sγPNA/BNP into intracranial tumors and captured the fluorescence signal of TAMRA-sγPNA (Fig. 6A). As expected, a strong fluorescence signal was detected in the brain using IVIS 3 h after CED (day 0), and brain cryosection images showed sγPNA/BNP accumulated mainly around the injection site after local delivery. After one day, we observed sγPNA/BNP distributed more widely from the injection site and spread throughout the tumor region in the brain, as indicated by Ki67, a marker for cell proliferation (Fig. S17). After 7 days, comparatively less fluorescence was detected in the brain, but a visible signal was still retained in the tumor region until day 14, suggesting that sγPNA molecules were well retained in the tumor for up to two weeks after CED (Fig. S18). Thus, encapsulation of sγPNA into BNP provided sustained retention within tumor which appears to be suitable for intracranial anti-miR treatment.

**Fig. 6.**
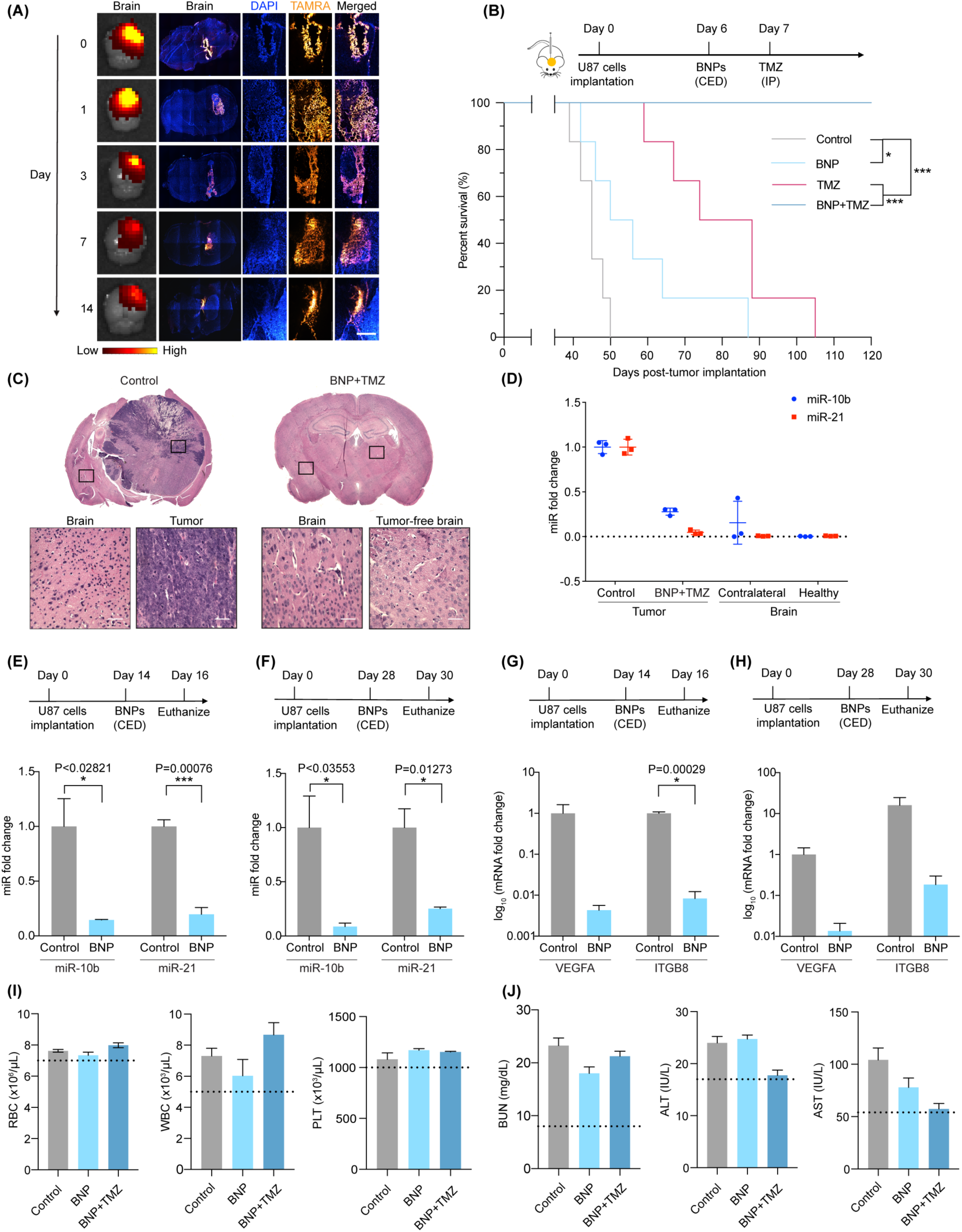
The combination of sγPNA/BNP and Temozolomide (TMZ) improves the survival in an orthotopic mice model of glioblastoma. **(A)** Biodistribution of BNP containing sγPNA-TAMRA on day 0, 1, 3, 7 and 14 after convention enhanced delivery (CED) via IVIS and microscopic imaging. Scale bar, 200 µm. **(B)** Survival of mice bearing U87 derived intracranial gliomas after treatment with sγPNA/BNP, TMZ and combination treatment of sγPNA/BNP with TMZ. NPs were administered by CED at 200 mg/ml dose and TMZ was administered intraperitoneally at 25 mg/kg dose. **(C)** H&E staining of control and sγPNA/BNP+TMZ treated mice brain at the end of the survival study. Scale bar, 50 µm. **(D)** The levels of miR-21 and miR-10b in gliomas of control and sγPNA/BNP +TMZ treated mice at the end of survival study. The levels of miR-10 and miR-21 were at similar levels as contralateral (to injection site) brain region and healthy mice brain. Results are represented as mean ± SD (n=3 biologically independent animals). **(E)** The gene expression levels of miR-10b and miR-21 in mice gliomas 48 h after CED of sγPNA/BNP on day 14. **(F)** The gene expression levels of miR-10b and miR-21 in mice gliomas 48 h after CED of sγPNA/BNP on day 28. **(G)** The levels of VEGFA and ITGB8 in tumors 48 h post CED on day 14. **(H)** The levels of VEGFA and ITGB8 in tumors 48 h post CED on day 28. Results are represented as mean ± SD (n=3 biologically independent animals). **(I)** Clinical chemistry of blood samples including white blood count (WBC), platelets (PLT), red blood cells (RBCs) from mice after 48 h of administration of sγPNA/BNP and sγPNA/BNP+TMZ. **(J)** Blood biochemistry including blood urea nitrogen (BUN), alanine aminotransferase (ALT), and aspartate transaminase (AST) from mice after 48 h of administration of sγPNA/BNP and sγPNA/BNP+TMZ. Results are represented as mean ± SD (n=3 biologically independent animals). sγPNA/BNP are physical mixture of sγPNA-21 BNP and sγPNA-10b BNP. BNP indicates PLA-HPG-CHO nanoparticles.

### Improved survival in orthotopic GBM tumor model

The significant tumor retention provided us a strong rational to evaluate the therapeutic efficacy of the sγPNA/BNP *in vivo*. Intracranial U87 tumors were generated in immunocompromised mice and NPs were administered via CED 6 days post tumor inoculation (Fig. 6B). The combination of sγPNA/BNP and TMZ was also evaluated by administering a single intraperitoneal injection of TMZ one day after CED infusion. In animals receiving sγPNA/BNP, median survival was significantly increased to 53 days compared to the untreated control groups (45 days), confirming the oncomiR inhibition mediated by sγPNAs effectively delayed tumor growth. Combination treatment of sγPNA/BNP plus TMZ (25 mg/kg) greatly improved survival and all the animals in this group (n=6) survived over 120 days. At the end of the study, the animals receiving the combination treatment (TMZ plus BNP) appeared to be tumor-free while control animals were found to be hunched due to tumor burden and neurological decline (Supplementary Video 1 and 2). Administration of TMZ (25 mg/kg) prolonged the median survival to 81 days, confirming therapeutic benefit, but was less effective than the combination with sγPNA/BNP. Further, animals treated by lower doses of TMZ (12.5 mg/kg) did not provide any significant improvement (p=0.0853) in survival comparing to the control group (Fig. S19), while the same dose of TMZ combined with sγPNA/BNP extended survival time over 100 days. These results indicated that sγPNA/BNP in combination with TMZ produced the longest survival time by successfully suppressing tumor growth and reducing the drug dose. Analysis of H&E and Ki67 brain sections of one survivor that received combination treatment and one untreated control animal clearly showed the presence of a fully developed tumor localized within the brain of the untreated mouse (Fig. 6C, S20, left), while the animal in sγPNA/BNP combination treatment group showed no evidence of tumor in the brain (Fig. 6C, S20, right).

### *In vivo* knockdown of miR-10b and miR-21

At the end of tumor survival study, we measured the relative levels of miR-10b and miR-21 in the tumors of untreated control animals and brain tissues around the injection site of sγPNA/BNP plus TMZ treatment group (Fig. 6D). Compared to control group, animals in the combined treatment group showed 72% knockdown of miR-10b and 95% decrease in miR-21 expression in the brain tissues ipsilateral to the injection site, indicating successful oncomiR inhibition mediated by sγPNA/BNP plus TMZ. The expression level of miR-10b and miR-21 in the contralateral hemisphere and healthy mice brains were also measured as comparisons.

To further investigate the role of sγPNA/BNP in oncomiR inhibition, we performed two separate experiments where sγPNA/BNP were administered 14 d post tumor implantation or 28 d post tumor implantation following the treatment schedule indicated in Fig. 6E and 6F, respectively. Tissues at the tumor site were harvested to examine expression levels of oncomiRs. qPCR results revealed that sγPNA/BNP mediated treatment resulted in statistically significant downregulation of miR-10b and miR-21 in tumors at different growth stages, further confirming efficient suppression of both targets. In addition, the levels of VEGFA and ITGB8 in the sγPNA BNP treated animals were evaluated; a consistent downregulation was found compared to control animals (Fig. 6G, 6H).

### Toxicity assessment of sγPNA/BNP and TMZ

We also assessed the toxicity of different treatments by performing complete blood counts, serum biochemistry, and histopathological analysis. As shown in Fig. 6I and S21, sγPNA/BNP alone and in combination with TMZ did not alter white blood cell (WBC), platelet (PLT), red blood cell (RBC) or other blood components when compared with control animals. No significant difference was found in liver enzymes (ALT, AST) and renal function markers (BUN) in mice after receiving treatments (Fig. 6J, S22). An independently conducted pathological analysis suggested that no obvious tissue changes were found in the treated animals when compared to control animals upon examination of the H&E stained major organs (Fig. S23). Overall, these data confirmed the safety of sγPNA/BNP in combination with TMZ.

### Improved survival in patient derived xenograft (PDX) mice model

We next tested the efficacy and safety of this combination strategy on a patient-derived GBM (G22) cells and found superior cellular uptake of sγPNA-loaded BNP *in vitro* (Fig. 7A, S24A). Significant downregulation of miR-10b and miR-21 and elevated response to TMZ treatment were also observed on PDX cells (Fig. S24B and C). Patient-derived orthotopic tumor model was established by injecting G22 cells intracranially in immunocompromised mice, as previously described (*60*), and treatments were performed as described above (Fig. 7B). Similarly, sγPNA/BNP combined with TMZ (25 mg/kg) greatly prolonged survival and 80% of animals survived over 120 days (one died on day 90, n=5). Animals receiving combination treatment that survived until the end of the study exhibited normal physical activity in comparison to untreated control animals, which had signs of morbidity as early as day 39 due to tumor development (Supplementary Video 3 and 4). Histological analysis of H&E and Ki67 stained sections further confirmed that brains of surviving BNP plus TMZ treated animals were tumor free (Fig. 7C, S25). Animals receiving sγPNA/BNP treatment or single dose of 25 mg/kg TMZ only extended median survival to 69 days and 79 days, respectively compared to the untreated controls (51 days), much less efficient than the combination of sγPNA/BNP with TMZ.

**Fig. 7.**
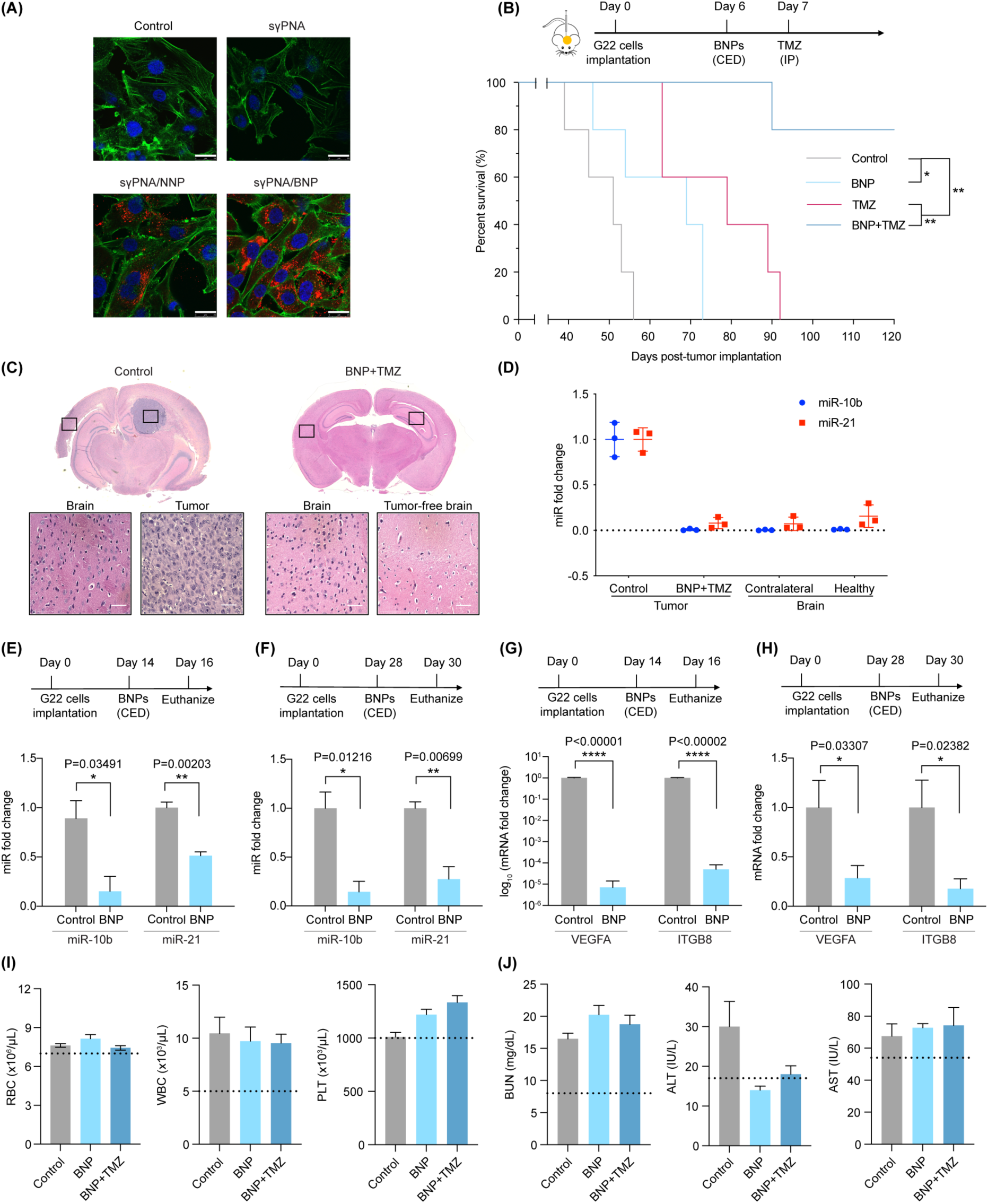
Efficacy of targeting oncomiR-10b and 21 via BNP delivered sγPNAs in patient derived xenograft (PDX) mice model of glioblastoma. **(A)** Confocal microscopic images of G22 cells treated by free sγPNA or sγPNA loaded NPs. PNAs were conjugated with TAMRA (red), F-actin was labelled with phalloidin (green) and nucleus was stained with Hoechst (blue). Scale bar, 25 µm. **(B)** Survival of mice bearing patient (G22) derived intracranial gliomas after treatment with BNPs containing sγPNA, TMZ and combination treatment of BNP with TMZ. BNPs were administered by convention enhanced delivery (CED) at 200 mg/ml dose and TMZ was administered intraperitoneally at 25 mg/kg dose. **(C)** The histology of H&E-stained control and BNP+TMZ treated mice brain at the end of the survival study. Scale bar represents 75 µm. **(D)** The levels of miR-21 and miR-10b in gliomas of control and BNP+TMZ treated mice at end of survival study. The levels of miR-10 and miR-21 were at similar levels as contralateral (to injection site) brain region and healthy mice brain. Results are represented as mean ± SD (n=3 biologically independent animals). **(E)** The gene expression levels of miR-10b and miR-21 in mice gliomas 48 h after CED of NPs on day 14 and on day 28 post tumor implantation. Results are represented as mean ± sem (n=3 biologically independent animals). **(F)** The levels of downstream targets VEGFA and ITGB8 in mice gliomas 48 h after CED of NPs on day 14 and day 28. Results are represented as mean ± sem (n=3 biologically independent animals). **(H)** Clinical chemistry of blood samples including white blood count (WBC), platelets (PLT), red blood cells (RBCs) from mice after 48 h of administration of BNP and BNP+TMZ. **(I)** Blood biochemistry including blood urea nitrogen (BUN), alanine aminotransferase (ALT), and aspartate transaminase (AST) from mice after 48 h of administration of BNP and BNP+TMZ. Results are represented as mean ± sem (n=3 biologically independent animals). sγPNA/BNP are physical mixture of sγPNA-21 BNP and sγPNA-10b BNP. NNP indicates PLA-HPG nanoparticles and BNP indicates PLA-HPG-CHO nanoparticles.

*In vivo* knockdown efficiency was also assessed on PDX model. The relative levels of miR-10b and miR-21 at the end of survival study were found low and comparable to those in the contralateral hemisphere and healthy mice brains (Fig. 7D). 99% knockdown of miR-10b and 92% knockdown of miR-21 were observed in the animals treated by sγPNA/BNP combined with TMZ. Significant downregulation of miR-10b and miR-21 in the tissue at tumor site was observed when administration of sγPNA/BNP was performed 14 days after tumor implantation or 28 days after tumor implantation (Fig. 7E, F). Similarly, the expression levels of VEGFA and ITGB8 were significantly lower than the untreated controls (Fig. 7G, H). Collectively, we found that sγPNA-loaded BNP resulted in successful inhibition of miR-10b and miR-21 and substantial improvement in animal survival in orthotopic PDX animals, consistent with the findings on intracranial U87 tumors.

Complete blood counts, serum biochemistry, and histopathological analysis were conducted to evaluate toxicity. Fig. 7I-J and S26-27 showed no significant difference between the treated animals and controls in terms of blood components including WBC, PLT, RBC and liver enzymes (ALT, AST) and renal function markers (BUN). H&E-stained sections of multiple organs, such as liver, spleen, kidney, cardiac muscle and lung showed no significant differences between treated animals and untreated controls in terms of inflammation or cell death (Fig. S28).

## DISCUSSION

Despite significant attention in preclinical and clinical research, GBM remains an aggressive disease, with limited survival and poor treatment options (*61, 62*). Little progress has been made toward improved survival outcomes in GBM patients over the standard of care, which includes surgical resection, radiation therapy plus TMZ (*63, 64*). Reasons for this failure include the lack of powerful therapeutic agents, restricted entry of drugs into intracranial tumors due to blood-brain-barrier (BBB), and tumor heterogeneity driven by multiple coordinated signaling pathways. All of these issues present challenges for GBM therapy (*65*). Here, we sought to investigate a potential solution that would involve an addition to the standard of care: infusion of highly effective sγPNA anti-miRs directly into tissue harboring tumor cells, with the sγPNAs packaged into NPs that are taken up in tumor cells. This strategy was targeted to two GBM-specific oncomiRs to overcome intrinsic resistance to the induction of cell death.

Several chemically modified oligonucleotides (antimiRs) exhibiting enzymatic stability and high binding affinity have been explored for targeting oncomiRs, including locked nucleic acid (*66*), 2-O methyl oligonucleotides (*16*), morpholinos (*67*), and PNAs (*32*). Based on their metabolic stability and binding affinity, PNA oligomers are efficient agents to inhibit the function of microRNAs (*68*). Modifications of PNAs at the γ-position can address many of the shortcomings of conventional PNAs, further maximizing antagonizing effect on the function of target microRNA (*69*). Here, we designed short serine-γPNAs conjugated with cationic arginine residues. Gel-shift assays showed that the synthesized sγPNAs (sγPNA-21, sγPNA-10b) bind with miR-10b and miR-21 separately, indicating specific and strong affinity for target oncomiRs. The sγPNAs encapsulated in BNP exhibited a greater miRNA inhibition effect in comparison with non-modified full length PNAs loaded in the same carrier. These short γPNA oligomers have enhanced hybridization due to their γ-modification and cationic residues, and we predicted that they would improve therapeutic efficacy.

In addition to efficacy, delivery of next generation antimiRs to the CNS remains an enormous challenge that need to be resolved. The efficacy of GBM therapies is significantly limited by the presence of blood-brain-barrier (*70*), however local delivery approaches have been beneficial to achieve intracranial distribution. Unlike diffusion-based methods (*71*), CED utilizes positive pressure flow and a catheter to achieve stereotactic placement and direct infusion of drugs into the tumor resection cavity providing large distribution volumes (*72*). In addition to intracranial delivery, high interstitial distribution can be achieved by adjusting the flow, distribution can be monitored in real time, and anti-tumor efficacy can be achieved at low doses, hence minimizing neurotoxicity as well as systemic toxicity (*73*). CED has been employed in multiple clinical trials for intracranial delivery of chemotherapeutics (*74*), antisense oligonucleotides (*75*), and liposomal vectors for gene therapy (*76*). In addition, CED has been investigated at non-clinical and pre-clinical stages for delivery of NPs (*77*) as well as viral vectors (*78*) for treatment of gliomas. Of note, recent clinical trials suggest that CED is safe and feasible but the present therapeutic approaches fail to significantly improve survival of GBM (*46*). This is most likely due to multiple reasons such as technical factors (*79*) (regions of the brain, catheter placement, rate of infusion), but more importantly, most drugs have short brain half-lives and are eliminated quickly after the infusion stops. We believe that infusion of NPs can improve current CED strategies (*80*) as NPs offer improved brain retention and sustained drug release for days to weeks after the end of infusion.

To achieve efficient cellular delivery, the sγPNAs were encapsulated in PLA-HPG nanoparticles. We previously demonstrated that nanoparticles composed of PLA-HPG (NNP), and a bioadhesive version, PLA-HPG-CHO (BNP), were safe and advantageous for delivery of PNAs to the brain (*26, 52*). Here, we observed 2- and 4-fold increase in cellular uptake with sγPNA loaded NPs formed from NNP and BNP respectively, over free sγPNAs. The preferential association of BNP with tumor cells also resulted in an enhanced oncomiR suppression *in vitro* analyzed by qRT-PCR. The expression of both miR-10b and miR-21 were significantly downregulated after sγPNA NP treatment. We hypothesize that altered miRNA levels in tumor cells might affect the expression of gene products that stimulate cell proliferation or that induce apoptosis, blocking tumor cells from developing into a proliferative state.

High throughout profiling revealed the overexpression of miR-10b and miR-21 in a large portion of human gliomas (*81*). Based on the specific molecular aberrations in GBM tumors, we designed sγPNAs targeting these two oncomiRs and prepared formulations loaded with sγPNA oligomers. Upon simultaneous inhibition of both targeted oncomiRs *in vitro*, we observed effective cell apoptosis in glioma cells treated by sγPNA/NP. A majority (54%) of apoptotic-associated cell death was achieved by exposure to 2 mg/mL sγPNA/BNP. These results suggested modest anticancer activity mediated by miRNA inhibition in tumor cells. We noticed that TMZ combined with sγPNA NPs induced a larger degree of cell apoptosis than TMZ or NPs alone. Therefore, we evaluated the cell killing activity of this combination strategy *via* cell survival assays *in vitro*, speculating improved chemosensitivity in treated tumor cells. As expected, combination with sγPNA/BNP effectively promoted tumor cell responses to TMZ, with significantly reduced cell viability than TMZ alone. Furthermore, this sensitization effect to chemotherapy mediated by γPNA/NP acted in a synergistic manner, as analyzed by the additive model (observed/expected ratio=0.644, Table S2) (*82, 83*). Tumor cell-specific cytotoxicity was also observed in this combination treatment, which displayed limited toxicity against healthy cells.

Limited therapeutic options, the presence of BBB and lack of progress in systemic delivery motivated more direct approaches for GBM therapy, such as local delivery via CED (*84, 85*). Wide distribution of truly effective agents by CED are expected to influence the overall therapeutic outcomes. With this in mind, CED of sγPNA encapsulated BNP was utilized to improve intracranial drug distribution and prolong the survival of tumor-bearing animals. We observed up to two-week retention and widespread distribution of sγPNA BNP within the intracranial tumor after CED infusion. This is consistent with our recently reported results where BNPs led to persistent presence of encapsulated camptothecin (∼50%) in the squamous cell carcinoma tumor at 10-day post tumor injection (*50*). We attribute the dramatically improved tumor retention to the enhanced association with tumor cells provided by the bioadhesive surface modifications of aldehyde-rich BNP. Based on our *in vitro* and *in vivo* results, we believe BNP-mediated increased tumor cell internalization and sustained drug release bring advantages to localized drug delivery and GBM therapy, particularly for invasive clinical approaches. Moreover, longer tumor retention is critically important when nucleic acid therapies are combined with other approaches (such as chemotherapy, radiotherapy) which require multiple doses over time.

In our survival study, CED of sγPNA-loaded BNP plus a single dose of TMZ significantly prolonged survival time of animals bearing intracranial U87 tumors and patient-derived G22 tumors. Although each of the monotherapies, sγPNA/BNP and TMZ alone showed some effect in delaying tumor progression and extending median survival, neither of them was able to completely inhibit tumor growth partially explaining the current limited efficacy of single targeted GBM therapy. Here we showed that all the U87 tumor-bearing animals receiving sγPNA/BNP plus TMZ treatment were tumor-free long survivors (>120 days); this robust antitumor activity could be beneficial for lowering the drug dose. For example, sγPNA/BNP combined with a reduced dose of TMZ (12.5 mg/kg) successfully increased survival time of tumor-bearing animals to over 100 days. Similar therapeutic benefits were reproduced on patient-derived GBM tumors thus leading to significantly improved animal survival (P<0.0018, vs control). We hypothesize that suppression of key oncotargets (miR-10b, miR-21) by anti-miR sγPNAs enhanced the sensitivity of glioblastoma cells to chemotherapy, resulting in an elevated response to TMZ treatment.

In conclusion, we propose a novel therapeutic approach to deliver anti-miR sγPNA-loaded BNP with bioadhesive surface modifications via CED to intracranial glioblastoma tumors. Two oncomiRs, miR-10b and miR-21, were targeted simultaneously, resulting in cooperative oncomiR inhibition and sensitization of tumors to TMZ treatment. Combined anti-miR sγPNA NPs with TMZ resulted in a remarkable delay of tumor growth and prevention of disease relapse. Thus, our strategy based on available clinical approach may improve eventual glioblastoma therapeutic outcome.

## MATERIAL AND METHODS

### MATERIAL

Poly (lactic acid) (Mw=20.2 kDa, Mn=12.4 kDa) was purchased from Lactel. Ethyl acetate, acetonitrile and DMSO were obtained from J.T. Baker. Temozolomide (TMZ) was obtained from Enzo Life Sciences. Human glioblastoma cell lines U87 and LN-229 was obtained from ATCC, human astrocyte was kindly provided by Dr. Ranjit Bindra at Yale. G22 (patient-derived xenograft (PDX) cells) were obtained from Jann Sarkaria (Mayo Clinic, Rochester, MN). The cells were grown in DMEM medium (Invitrogen) supplemented with 10% fetal FBS, 1% Pen-Strep and cultured at 37 °C with 5% CO2 in a humidified chamber.

### METHODS

#### Synthesis of PNA Oligomers

PNAs were synthesized via solid phase synthesis using MBHA (4-Methylbenzhydrylamine) resin and standard Boc chemistry procedures as reported previously (*42*). Regular Boc-monomers and serine-γPNA-Boc-monomers (A, T, C, G) purchased from ASM Chemicals and Research (Germany) were used. Three arginine residues were conjugated to N terminus or 5’ end of PNAs. Carboxytetramethylrhodamine (TAMRA) dye, bought from VWR (Pennsylvania, USA), was further conjugated to 5’ end with Boc-MiniPEG-3 linker in between. After completion of synthesis, PNAs were cleaved from the resin using trifluoroacetic acid: trifluoromethanesulfonic acid: thioanisole: m-cresol at a ratio of 6:2:1:1 as cleavage cocktail and precipitated using diethyl ether. PNAs were further purified using RP-HPLC to obtain the pure fraction of PNA. The molecular weight of purified PNAs was confirmed using matrix-assisted laser desorption/ionization-time of flight (MALDI-TOF) spectrometry. The concentration of PNAs in water was determined using UV-Vis spectroscopy. The amount of PNA was then calculated using extinction coefficient of PNA obtained by combining the extinction coefficient of individual monomers of the sequence. **Gel shift assay**

The binding of PNAs with the target oncomiRs 10b and 21 was determined by incubating PNAs with target oncomiRs 10b and 21 in a buffer simulating physiological ionic conditions (10 mM NaPi, 150 mM KCl and 2 mM MgCl_2_) at 37°C for 16 hours in thermal cycler (Bio-Rad, USA). The samples were separated using 10% non-denaturing polyacrylamide gel and tris/boric acid/EDTA buffer (TBE) at 120V for 35 minutes. For detection of bound and unbound fraction of target oncomiRs, gels were stained using SYBR-Gold (Invitrogen, USA) followed by imaging in a Gel Doc EZ imager (Bio-Rad, USA).

#### Nanoparticle preparation

##### sγPNA/NNP

sγPNA loaded NNP were prepared using emulsion-evaporation method as previously reported. Fifty (50) mg polymer (PLA-HPG) was dissolved in 2.4 mL of ethyl acetate overnight. 50 nmol sγPNA was dissolved in 0.6 mL of DMSO and then added to polymer solution, obtaining PNA/polymer mixture. The resulting solution was added dropwise to 2 mL of DI water under a strong vortex, and then sonicated 10 s for 4 cycles. The emulsion was diluted in 20 mL deionized (DI) water and concentrated in a rotovap for 20 min. The particle solution was transferred to an Amicon centrifugal filter unit (100 KDa) and washed twice by water. Lastly, the obtained particles were resuspended in DI water and snap-frozen in aliquots. sγPNA-21 loaded NNP and sγPNA-10b loaded NNP were mixed at 1:1 molar ratio to obtain sγPNA/NNP before further use. Scr-sγPNA and regular PNA loaded NNP were prepared using the same method.

##### sγPNA/BNP

To create sγPNA/BNP, 25 mg/mL sγPNA/NNP (as above) was incubated with 0.1 M NaIO_4_ (aq) and 10x PBS (1: 1: 1, v: v) for 20 min on ice. 0.2 M NaSO_3_ at 1:1 vol ratio was added to quench the reaction. The particle solution was washed with water at 13,000 rcf using a centrifugal filter unit (Amicon, 100 KDa) and resuspended in DI water. Regular PNA loaded BNPs were synthesized with the same method.

#### Nanoparticle characterization

##### Transmission electron microscopy (TEM)

For TEM imaging, 2 µL of particle solution (20 mg/mL) was applied on a CF400-CU grid (Electron Microscopy Sciences) for 1 min. Extra liquid was carefully removed and the grid was stained by one drop NANO-W (Nanoprobe) for 1 min. Liquid was removed and sample was air dried before imaging. Images were obtained using Tecnai Osiris (FEI).

##### Size and ζ potential

The hydrodynamic diameter of NPs was measured by dynamic light scattering (DLS) using a Malvern Nano-ZS (Malvern Instruments). NPs were diluted to 0.2 mg/mL with DI water before measurement. The same particle solution was loaded into a disposable capillary cell to measure ζ potential on the Malvern Nano-ZS.

##### Size stability in aCSF

Particle solutions were incubated in artificial cerebrospinal fluid (aCSF, Harvard Apparatus) at 37 °C and measured by DLS at designated time points.

##### sγPNA loading and release

100 µL of particle solution was lyophilized in a pre-weighed tube to measure NP yield. Following lyophilization, NPs were dissolved in acetonitrile and incubated for 24 h at room temperature. Absorbance at 260 nm was read by a Nanodrop 8000 (Thermo Fisher) to measure sγPNA loaded in the NPs. Scr-sγPNA and regular PNA loading efficiencies were determined using the same method. Release profile of sγPNA from different NP formulations was analyzed by incubating 10 mg NPs in 1 mL PBS (pH 7.4) in a shaking incubator at 37 °C. At predetermined time points, aliquots were taken out and centrifuged using Amicon centrifugal filter unit (100 KDa). Filtrates were collected for analysis.

#### Cellular uptake of sγPNA loaded nanoparticle

Cells were seeded in 24-well plates at a density of 50,000 cells/well and incubated with either 0.5 mg/mL NP or same concentration of free sγPNA for 24 h. sγPNA oligomers used in this experiment were labeled with TAMRA for fluorescent evaluation. The cells were harvested, and cell uptake was determined from TAMRA fluorescence per cell using Attune NxT (Invitrogen) flow cytometer and FlowJo software for data analysis. For microscopic observation, U87 cells were cultured in a 20 mm glass-bottom dish (20,000 cells/dish) before treatments. Cells were exposed to NPs (1 mg/mL) or free sγPNA for 24 h and washed by PBS after treatments. After 4% paraformaldehyde fixation, cells were stained with Alexa Fluor 488 phalloidin (Life Technologies) and DAPI, and observed by SP5 confocal microscope (Leica).

#### Quantitative real-time polymerase chain reaction (qRT-PCR)

The knockdown of miR-10b and miR-21 were analyzed by qRT-PCR. Cells were seeded in 24-well plates at a density of 200,000 cells/well. Cells were treated with various formulations at a PNA concentration of 300 nM. Scr-sγPNA loaded NPs and regular PNA loaded NPs with the same total PNA concentration were applied as control groups. After 72 h, total RNA was extracted using the mirVana miRNA Isolation Kit (Ambion). cDNA synthesis was performed using TaqMan Advanced miRNA cDNA Synthesis Kit (Thermo Fisher). PCR reactions were performed with TaqMan Fast Advanced Master Mix (Thermo Fisher), and TaqMan Advanced miRNA Assays (Thermo Fisher) for miR-10b, miR-21 and miR-26b analysis. MiRNA levels were quantified using CFX Connect Real-Time PCR Detection System and CFX Manager Software (Bio-Rad). Relative expression was calculated according to the comparative threshold cycle (Ct) method and normalized by miR-26b. For evaluation of PTEN mRNA level, PCR reactions were performed using PTEN and GAPDH TaqMan Gene Expression Assays (Thermo Fisher) and quantified with CFX Real-Time PCR Detection System and CFX Manager Software. The results were calculated with Ct method and normalized by GAPDH.

#### Annexin V assay

Apoptosis was assessed using the FITC Annexin V Apoptosis Detection Kit (BD Pharmingen). Briefly, tumor cells were plated in 24-well plates at a density of 200,000 cells/well. The cells were treated with free sγPNA, sγPNA loaded NPs and/or TMZ for 72 h (200 nM sγPNA-21, 200 nM sγPNA-10b, 40 µM TMZ). The FITC Annexin V apoptosis detection was performed using flow cytometry in accordance with the manufacturer’s protocol, and the data were processed using FlowJo. FITC-Annexin V positive and PI-negative cell populations were identified as early apoptotic cells. Cells that stain positive for both FITC Annexin V and PI are either in the end stage of apoptosis or are undergoing necrosis. Cells that stain negative for both dyes are identified as alive.

#### Caspase 3/7 activity evaluation

U87 cells were plated in 96-well plates at a density of 5,000 cells/well and treated with various sγPNA nanoparticle formulations and TMZ (200 nM sγPNA-21, 200 nM sγPNA-10b, 40 µM TMZ). Treatments were removed 48 or 72 h later and enzymatic activities of caspase-3 and -7 were measured using Caspase-Glo 3/7 Assay (Promega) and read by a microplate reader (SpectraMax M5).

#### Cell viability assay

Cells were seeded in 96-well plates at a density of 5,000 cells/well and treated with increasing concentrations of NP formulations. Cell viability was evaluated by CellTiter-Glo Luminescent Cell Viability Assay (Promega) after 48 h and 72 h treatment. Luminescence was measured using a plate reader (SpectraMax M5) and relative cell viability was normalized to the viability of untreated cells. For TMZ involved combination studies, cells were seeded in 96-well plates at a density of 1,000 cells/well and exposed to sγPNA/NNP and sγPNA/BNP. After 72 h of treatments, NPs were removed and TMZ was added to the wells. Cell viability was measured as described above after 6 days of treatment.

#### RNA-sequencing

U87 cells were seeded in 24-well plates at a density of 50,000 cells/well and cultured overnight before use. sγPNA-21/BNP (150 nM sγPNA-21), sγPNA-10b/BNP (150 nM sγPNA-10b), sγPNA/BNP (150 nM sγPNA-21, 150 nM sγPNA-10b) were added to cells and incubated for 72 h. Total RNA from each sample was extracted using the mirVana miRNA Isolation Kit. The libraries were made using Illumina TruSeq Stranded mRNA library preparation. The sequencing was done using Illumina NextSeq 500.

#### Analysis of RNA-sequencing data and miRNA targets

Total counts per gene were quantified and used in further analysis. All downstream analyses were accomplished by R (3.6.3). Differentially expressed genes (DEGs) between different groups were identified by the package DESeq2 (1.26.0) with a filtering criteria of fold change (FC)≥1.5 and adjust p value (padj)<0.05 (*86*). The package cluster profiler (3.14.3) was used to identify specific pathways overrepresented in the DEGs, and significant pathways were picked out by setting p value cutoff =0.05 and q value cutoff =0.05 (*87*). Because the miRTarBase2020 is the updated version of the experimentally validated microRNA–target interaction database (*88*), the targets of miR10b-5p and miR21-5p were downloaded from the website (http://miRTarBase.cuhk.edu.cn/) and were used in the study.

#### TCGA GBM data analysis

The TCGA miRNA expression level-3 data and metadata containing survival information for GBM patients were downloaded from http://gdac.broadinstitute.org/. We ranked the GBM patients from high to low according to their miR 10b or miR 21 expression level, then labeled the top 25% patients as the miR-higher group, bottom 25% ones as the miR-lower group. One GBM patient would be marked as miR 10b & miR 21-higher when this patient was in both miR 10b-higher group and miR 21-higher group, and miR 10b & miR 21-lower patient was in both miR 10b-lower group and miR 21-lower group. Survival curves were performed by Kaplan–Meier analysis between miR higher and lower group, and were tested for significance using the Mantel-Cox log-rank test. A value of p<0.05 was considered statistically significant. Between miR 10b & miR 21-higher and miR 10b & miR 21-lower group, hazard ratio (HR) and confidence interval (CI) were also computed by function coxph in package survival (3.2-7).

#### *In vivo* study of sγPNA/BNP

All procedures were approved by the Yale University Institutional Animal Care and Use Committee (IACUC) and performed in accordance with the guidelines and policies of the Yale Animal Resource Center (YARC). Athymic nude mice (Charles River Laboratories, 6-7 weeks) were used for animal study.

#### Orthotopic tumor inoculation

Animals were anaesthetized using a mixture of ketamine (100 mg/kg) and xylazine (10 mg/kg) via intraperitoneal injection. Anesthetized animals were then placed in a stereotaxic frame and sterilized the scalp with betadine and alcohol. To expose the coronal and sagittal sutures, a midline scalp incision was created, and a burr hole was drilled 2 mm lateral to the sagittal suture and 0.5 mm anterior to the bregma. U87 or G22 cells (3.5 x 10^5^) in 3 µL PBS were injected into the right stratum over 3 min using a 10-µL Hamilton syringe. The animal was left 5 min for tissue equilibration before and after infusion. When infusion was finished, the burr hole was filled with bone wax and skin was stapled and cleaned.

#### Convection-enhanced delivery of sγPNA/BNP in the tumor bearing brain

CED in tumor bearing mice was similar to tumor inoculation by reopening the burr hole used for tumor inoculation. A micro-infusion pump (World Precision Instruments) was used to infuse 6 µL of NPs at a rate of 0.5 µL/min.

#### Tumor retention of sγPNA/BNP

Intracranial CED of sγPNA/BNP was conducted 10 days after tumor implantation using the same procedure as previously described. On day 0, 1, 3, 7 and 14 after CED administration, mice were sacrificed and brains were harvested and imaged using Xenogen IVIS. The fluorescence from each brain was quantified by the instrument software. Then, the isolated brains were embedded in an OCT compound, cut into 15 microns frozen sections, stained with DAPI or H&E (hematoxylin and eosin) and observed using a EVOS microscope (FL Auto 2).

#### Survival study in the tumor bearing brain

Tumors grew for 6 days before the administration of treatment. Intracranial CED of sγPNA/BNP was conducted following the same surgical procedure as described. TMZ (25 mg/kg) in PBS was administered intraperitoneally on day 7. Animals were monitored daily and weighed every week. Animals were euthanized once they showed clinical symptoms of tumor progression or greater than 15% weight loss. Tumors and major organs were harvested and fixed in 4% paraformaldehyde and sectioned for histochemical analysis. Total RNA in the tumor and contralateral hemisphere was isolated using mirVana miRNA Isolation Kit and analyzed by real-time PCR detection system as previously described.

#### Evaluation of miR-10b and miR-21 inhibition in tumor

To assess *in vivo* knockdown effect, sγPNA/BNP were administered by CED 14 d or 28 d post tumor inoculation. Two days after CED, mice were sacrificed and brains were harvested for RT-qPCR analysis. Tumor tissue was separated from the adjacent normal brain areas of isolated brains. Total RNA was extracted from tumor tissue using the mirVana miRNA Isolation Kit (Ambion). MiRNA levels were quantified using CFX Connect Real-Time PCR Detection System and CFX Manager Software (Bio-Rad) as described.

#### Evaluation of *VEGFA* and *ITGB8*

The levels of *VEGFA* and *ITGB8* were quantified in total RNA samples extracted from *in vitro* and *in vivo* tumor samples. The cDNA was synthesized using a high capacity cDNA reverse transcription kit (Thermo Fisher). The mRNA levels of *VEGFA* and *ITGB8* were quantified using TaqMan gene expression assays (*VEGFA*: Hs00900055, *ITGB8*: Hs001744546) (Thermo Fisher) and TaqMan fast advanced master mix (Thermo Fisher) on CFX connect Real-Time PCR Detection system (Bio-Rad). GAPDH was used as reference and relative fold change was calculated using the Ct method.

#### Toxicity study

For evaluation of the toxicity, tumor-bearing mice were administered of sγPNA/BNP via CED 14 d post tumor inoculation. After 24 h, TMZ (25 mg/kg) was injected intraperitoneally. Forty-eight (48) h post CED, blood from the retro-orbital venous plexus of each mouse was collected in EDTA tubes. The whole blood was analyzed using Sysmex XP-300 hematological analyzer (Sysmex) to obtain the complete blood count. Plasma was isolated from blood samples via centrifugation at 4500 rpm and 4°C for 10 minutes. Plasma samples were then analyzed by Antech Diagnostics to obtain the blood biochemistry analysis including LDH, AST, ALT, creatinine, and BUN. Major organs (heart, liver, spleen, lung, kidney) were isolated and sectioned for H&E staining. Blinded histological analysis of the tissue was conducted by pathologist at Yale medical school.

#### Statistical analysis

Data are presented as mean ± SD or SEM. Statistical significance was performed with Prism software (GraphPad) using one-way ANOVA. *p* < 0.05 as the minimal level of significance.

## Acknowledgments

We thank Anisha Gupta for discussions and suggestions.

## Funding

This work was supported by Hood Foundation award, and grants from the National Institutes of Health (CA241194 to RB and CA149128 to WMS).

## Author contributions

Conceptualization: RB, WMS

Methodology: YW, SM, RB, WMS

Investigation: YW, SM, YX, YD, AH

Visualization: YW, SM, YX, YD, AH, RF, RSB, WMS, RB

Funding acquisition: RB, WMS

Writing: YW, SM, HWS, YX, YD, RF, AH, RSB, WMS, RB

## Competing interests

The authors declare that they have no competing interests.

## Data and materials availability

All data associated with the study is present within the paper or supplementary materials. RNA sequencing data is available at the accession number (pending from GEO) in Gene Expression Omnibus (https://www.ncbi.nlm.nih.gov/geo/).

## Graphical Abstract

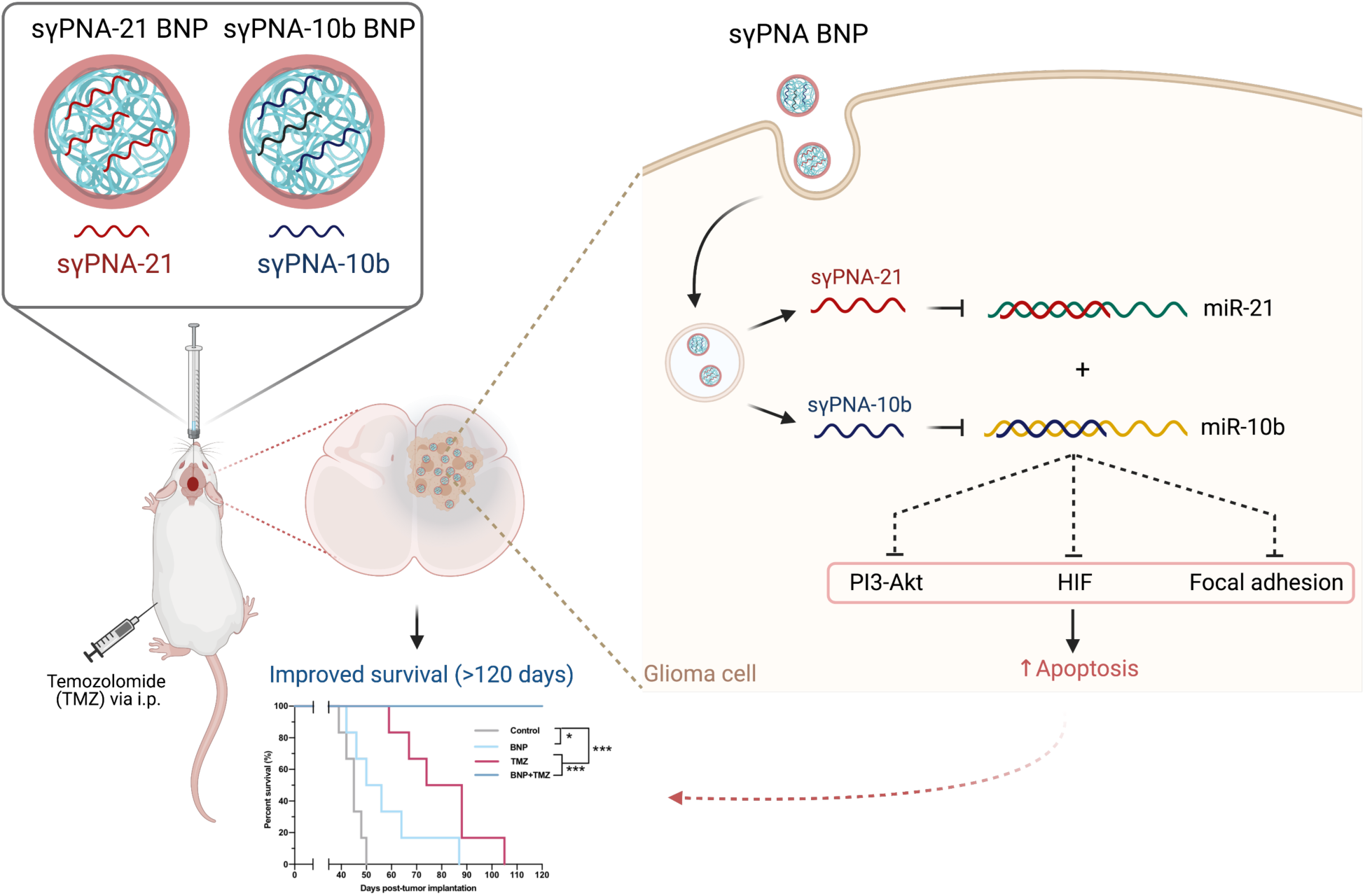

## Notes

### Competing Interest Statement

The authors have declared no competing interest.

