## Supplementary materials for "Anti-seed PNAs targeting multiple oncomiRs for brain tumor therapy"

**This file includes:**

**Supplementary figures**

Fig. S1. The HPLC chromatograms of the PNAs used in the study.

Fig. S2. Gel shift assay of PNAs with miR-21 and miR-10b.

Fig. S3. Gel-shift assay of sγPNA-21 and sγPNA-10b with the target miR 21 and miR 10b as well as mixture of both the target miRs.

Fig. S4. Quantification of bio-adhesiveness of sγPNA loaded NNPs and BNPs using poly-l-lysine (PLL) coated glass.

Fig. S5. Cellular uptake of sγPNA NNP/BNP in glioma cells.

Fig. S6. Gene expression of PTEN, miR-21, and miR-10b.

Fig. S7. Flow cytometry analysis of cell apoptosis after various treatments using Annexin V assay.

Fig. S8. sγPNA NNP/BNP induce apoptosis in U87 cells.

Fig. S9. RNA sequencing work flow.

Fig. S10. Differentially expressed genes after sγPNA BNP mediated knockdown of both miR-10b and miR-21.

Fig. S11. RNA sequencing analysis after sγPNA-21 BNP mediated knockdown of miR-21 in U87 cells.

Fig. S12. RNA sequencing analysis after sγPNA-10b BNP mediated knockdown of miR-10b in U87 cells.

Fig. S13. Heatmaps of PI3-Akt and focal adhesion pathway after miR-21 knockdown in U87 cells.

Fig. S14. Heatmaps of PI3-Akt pathway after miR-10b knockdown in U87 cells.

Fig. S15. The correlation of survival probabilities and levels of miR-10b or miR-21.

Fig. S16. The correlation of survival probabilities and levels of both miR-10b and miR-21.

Fig. S17. Immunostaining of Ki67 in TAMRA-sγPNA /BNP treated mice (CED) after one day.

Fig. S18. Mean fluorescence intensity of TAMRA-sγPNA in the brain.

Fig. S19. Survival study of animals treated with lower dose of TMZ (12.5 mg/kg) and combination treatment with TMZ and sγPNA/BNP.

Fig. S20. Ki67 staining of control and sγPNA/BNP+TMZ treated U87 tumor-bearing mice brain at the end of the survival study.

Fig. S21. *In vivo* toxicity analysis of BNP and BNP+Temozolomide (TMZ).

Fig. S22. The biochemistry analysis of blood samples from mice treated with BNP and BNP+TMZ after 48 hours of treatment.

Fig. S23. H&E staining of major organs including heart, liver, spleen, lung, liver, and kidney in all groups of mice from survival study.

Fig. S24. Cellular uptake and efficacy of sγPNA in G22 cells (patient derived glioblastoma cells).

Fig. S25. Ki67 staining of control and sγPNA/BNP+TMZ treated G22 tumor-bearing mice brain at the end of the survival study.

Fig. S26. The complete blood count analysis of blood samples from G22 PDX mice model treated with BNP and BNP+TMZ after 48 hours of treatment.

Fig. S27. The biochemistry analysis of blood samples from G22 PDX mice model treated with BNP and BNP+TMZ after 48 hours of treatment.

Fig. S28. H&E staining of major organs including heart, liver, spleen, lung, liver, and kidney in all groups of mice from survival study in G22 PDX mice model of glioblastoma.

**Supplementary tables**

Table S1. Molecular weight (MW) of perfect match PNAs used in the study.

Table S2. The synergistic effect of γPNA/BNP and TMZ combination treatment on U87 cells.

**Supplementary videos**

Supplementary video 1. Untreated mouse bearing U87 intracranial tumor on day 46 post tumor implantation.

Supplementary video 2. Mouse treated with BNP+TMZ (200 mg/mL BNP, 25 mg/kg TMZ) on day 120 after U87 tumor implantation.

Supplementary video 3. Untreated mouse bearing G22 (patient derived glioblastoma cells) intracranial tumor on day 39 post tumor implantation.

Supplementary video 4. Mouse treated with BNP+TMZ (200 mg/mL BNP, 25 mg/kg TMZ) on day 120 after G22 tumor implantation.

**Supplementary excels (RNA sequencing analysis)**

Supplementary Excel 1: RNAseq_raw_read_counts

Supplementary Excel 2: control vs pna-10b_DESeq2.results

Supplementary Excel 3: control vs pna-10b__log2FC_0.585_padj_0.05

Supplementary Excel 4: control vs pna-21_DESeq2.results

Supplementary Excel 5: control vs pna-21_log2FC_0.585_padj_0.05

Supplementary Excel 6: control vs pna-21+10b_DESeq2.results

Supplementary Excel 7: control vs pna-21+10b_log2FC_0.585_padj_0.05

Supplementary Excel 8: hsa-miR-10b-and-21 target genes

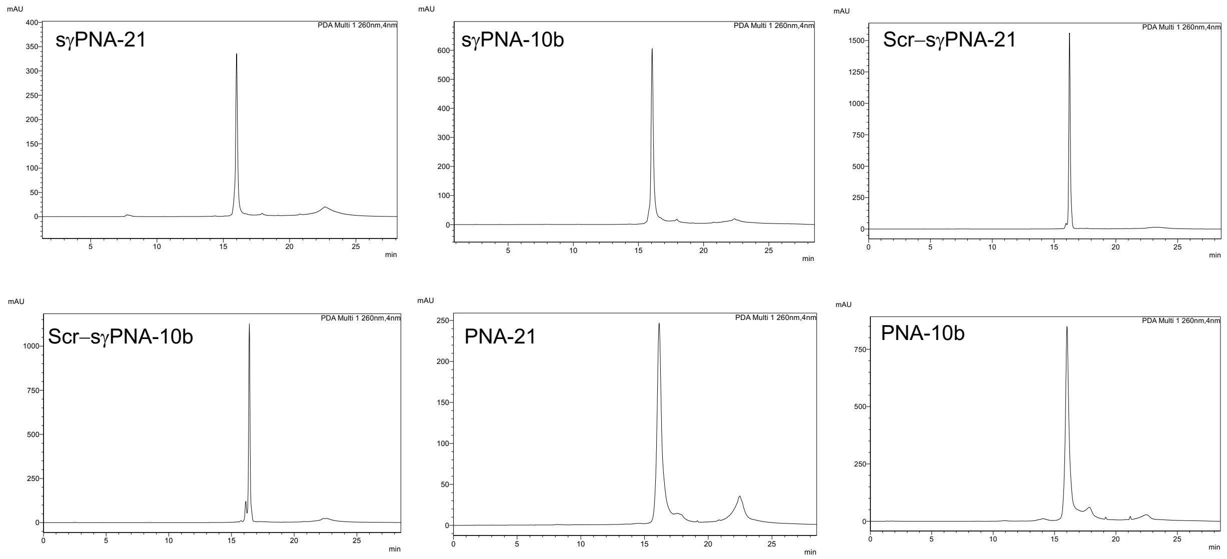


**Fig. S1. The HPLC chromatograms of the PNAs used in the study.**

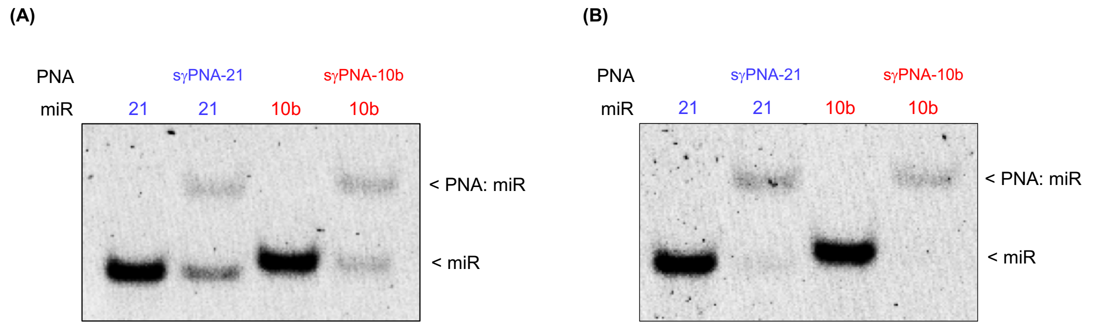


**Fig. S2. Gel shift assay of PNAs with miR-21 and miR-10b. (A)** Gel-shift assay of miR-21 and miR 10b targets with sγPNA-21 and sγPNA-10b, respectively at PNA: miR ratio of 2:1. **(B)** Gel-shift assay of miR-21 and miR 10b target with sγPNA-21 and sγPNA-10b, respectively at PNA: miR ratio of 4:1. PNAs were incubated with the target miRs in simulated physiological buffer conditions (10mM NaPi, 150 mM KCl and 2 mM $MgCl_2$) for 16 hours at 37°C followed by non-denaturing PAGE separation and SYBR Gold staining.

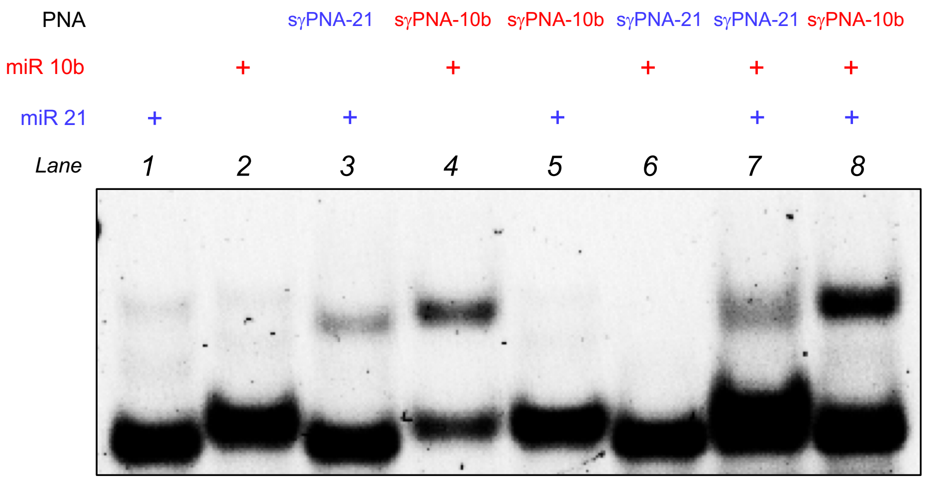


**Fig. S3. Gel-shift assay of sγPNA-21 and sγPNA-10b with the target miR 21 and miR 10b as well as mixture of both the target miRs**. PNAs were incubated with the target miRs at PNA: miR ratio of 1:1 in simulated physiological buffer conditions (10mM NaPi, 150 mM KCl and 2 mM $MgCl_2$) for 16 hours at 37°C followed by non-denaturing PAGE separation and SYBR Gold staining.

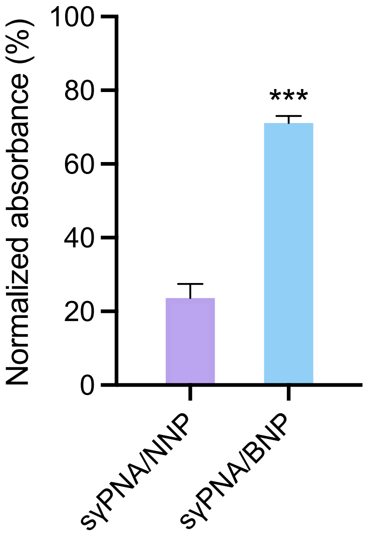


**Fig. S4. Quantification of bio-adhesiveness of sγPNA loaded NNPs and BNPs using poly-l-lysine (PLL) coated glass.** NPs were incubated with PLL glass cover slides for 30 min then washed with water. Acetonitrile was added to each slide to harvest the remaining sγPNA-loaded NPs. The concentration of remaining PNA on the slides was quantified using the methods described. Data was normalized to unwashed controls. ***$p < 0.001$ vs NNP.

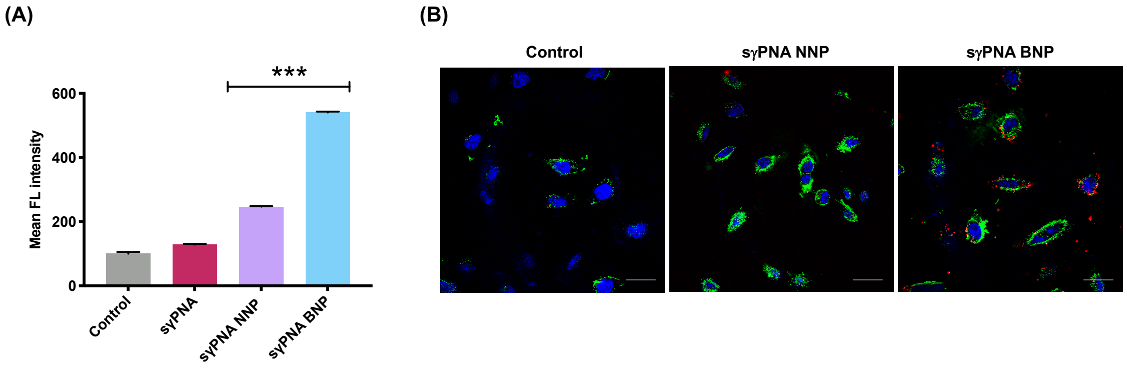


**Fig. S5. Cellular uptake of sγPNA NNP/BNP in glioma cells. (A)** Quantification of cellular uptake in U87 cells by flow cytometry analysis. ***$p < 0.001$. **(B)** Confocal microscopic images of LN229 cells treated by sγPNA loaded nanoparticles. PNAs were conjugated with TAMRA (red), F-actin was labelled with phalloidin (green) and nucleus was stained with Hoechst (blue). Scale bar, 20 μm.

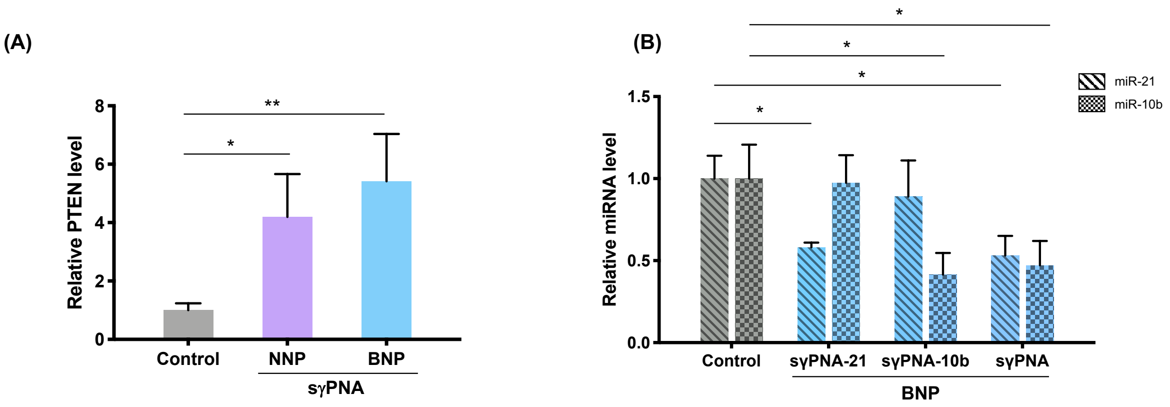


**Fig. S6. Gene expression of PTEN, miR-21, and miR-10b. (A)** Relative PTEN mRNA level in U87 cells transfected with sγPNA NNP and sγPNA BNP. ***$p < 0.001$ vs control, **$p < 0.01$ vs control and *$p < 0.05$ vs control. Data are expressed as mean ± SD (n = 3). **(B)** qRT-PCR analysis of relative miR-10b and miR-21 level in cells treated by sγPNA-21 BNP, sγPNA-10b BNP and sγPNA BNP nanoparticles. sγPNA BNP nanoparticles are physical mixture of sγPNA-21 BNP and sγPNA-10b BNP. NNP indicates PLA-HPG nanoparticles and BNP indicates PLA-HPG-CHO nanoparticles. **$p < 0.001$ vs control.

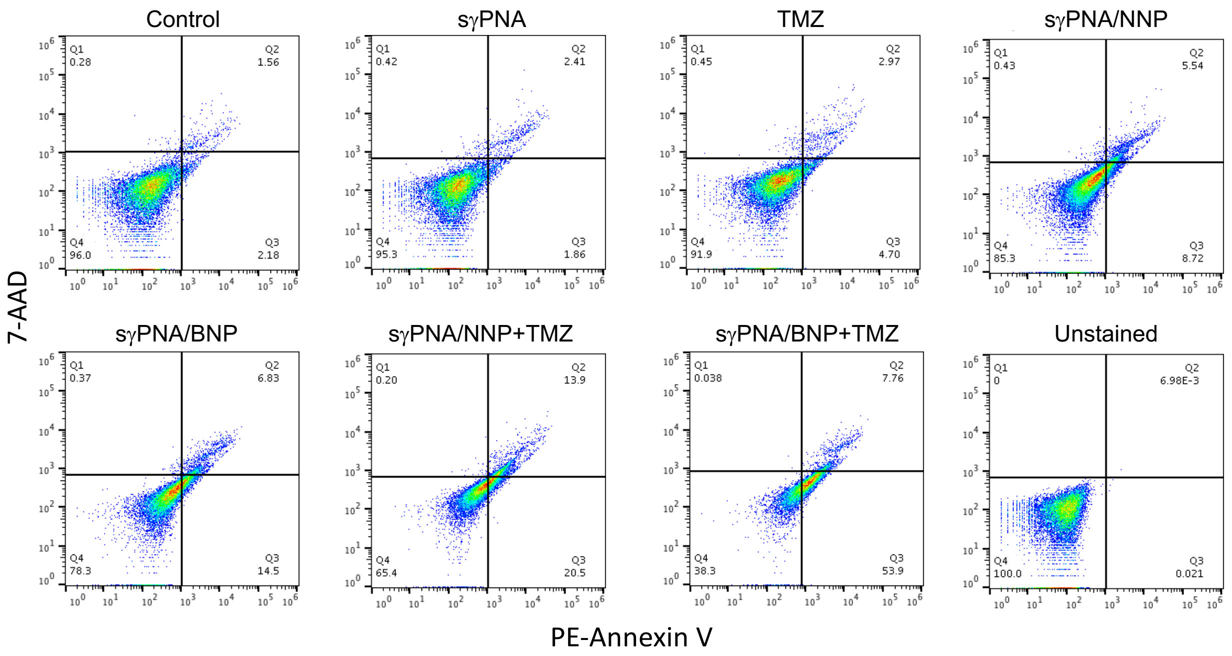


**Fig. S7. Flow cytometry analysis of cell apoptosis after various treatments using Annexin V assay.**

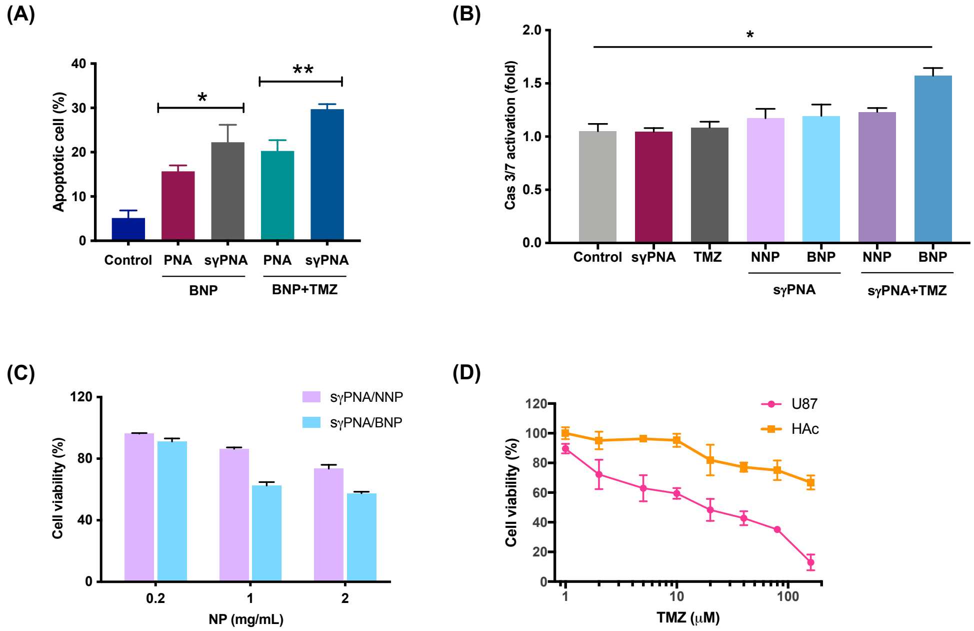


**Fig. S8. sγPNA NNP/BNP induce apoptosis in U87 cells. (A)** Quantification of apoptotic cells treated by PNA/BNP, sγPNA/BNP with and without TMZ by flow cytometry. $*p < 0.05$, $**p < 0.01$. **(B)** Caspase 3 and caspase 7 activities of cells treated by various formulations for 48 h. $*p < 0.05$. **(C)** Cell viability of U87 cells treated by increasing doses of sγPNA/NNP and sγPNA/BNP for 48 h. **(D)** Cell viability of tumor cells (U87) and human astrocytes (HAc) treated by sγPNA/BNP and increasing doses of TMZ for 72 h. sγPNA/BNP are physical mixture of sγPNA-21 BNP and sγPNA-10b BNP. PNA/BNP are physical mixture of PNA-21 BNP and PNA-10b BNP. NNP indicates PLA-HPG nanoparticles and BNP indicates PLA-HPG-CHO nanoparticles.

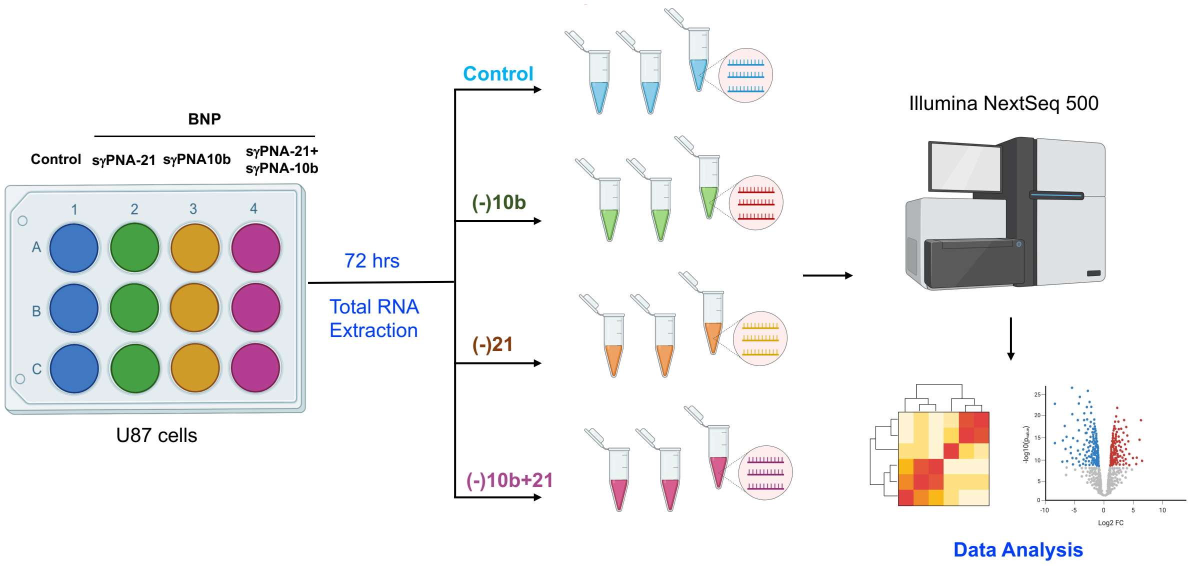


**Fig. S9. RNA sequencing work flow.** U87 cells were treated with BNP containing sγPNA-10b or sγPNA-21 or combination of BNP loaded with sγPNA-21 and sγPNA-10b for 72 hours. Total RNA was collected from the treated and untreated (control) cells for RNA sequencing analysis.

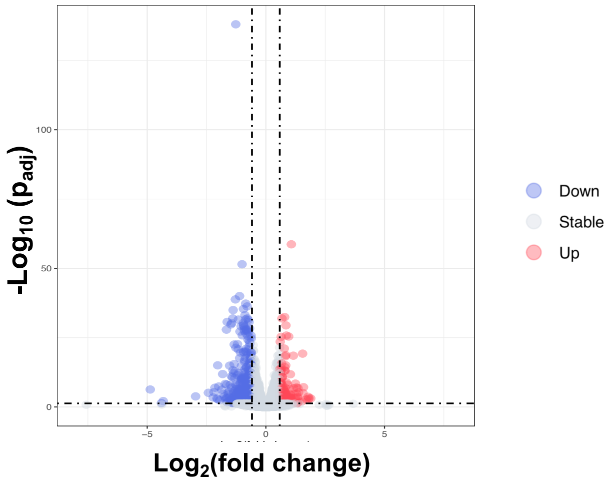


**Fig. S10. Differentially expressed genes after sγPNA BNP mediated knockdown of both miR-10b and miR-21.** The volcano plot showing differentially upregulated (red) and downregulated (blue) genes in U87 cells treated with sγPNA/BNP (physical mixture of sγPNA-21/BNP and sγPNA-10b/BNP) for 72 h when compared against the untreated cells (Control).

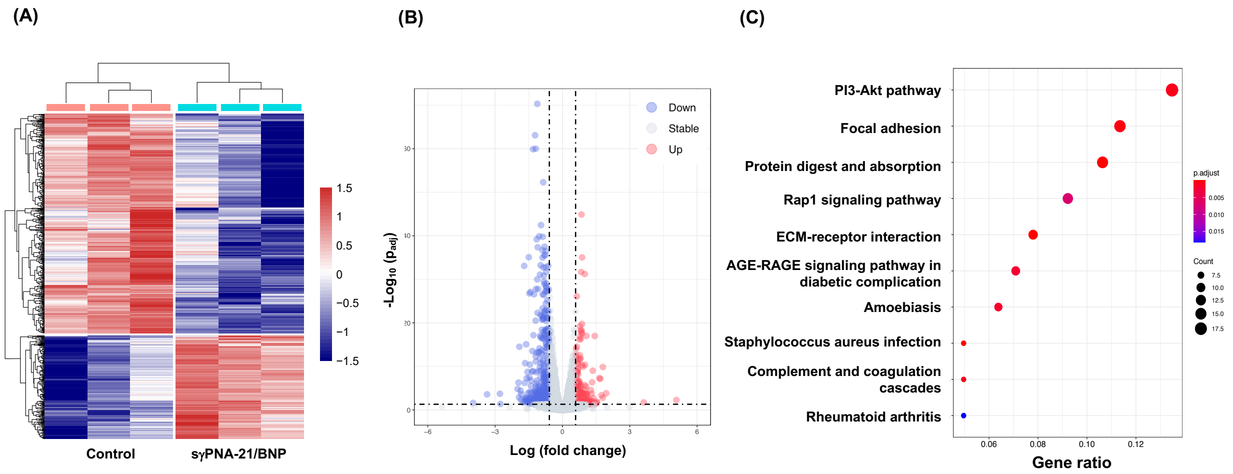


**Fig. S11. RNA sequencing analysis after sγPNA-21 BNP mediated knockdown of miR-21 in U87 cells. (A)** The heatmap showing hierarchical clustering of differentially expressed genes in U87 cells treated with sγPNA-21/BNP in comparison to the control (untreated). **(B)** The volcano plot showing differentially expressed genes in sγPNA-21/BNP treated U87 cells. **(C)** Gene ontology (GO) enrichment analysis of the differentially expressed downregulated genes in U87 cells treated with sγPNA-21/BNP in comparison to the control (untreated cells).

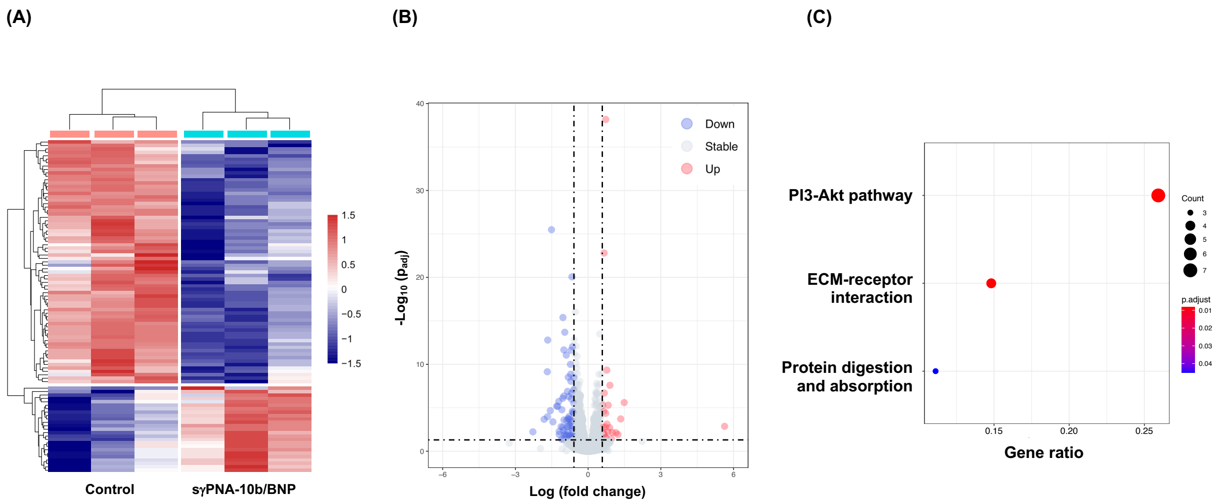


**Fig. S12. RNA sequencing analysis after sγPNA-10b BNP mediated knockdown of miR-10b in U87 cells. (A)** The heatmap showing hierarchical clustering of differentially expressed genes in U87 cells treated with sγPNA-10b/BNP in comparison to the control (untreated). **(B)** The volcano plot showing differentially expressed genes in sγPNA-10b/BNP treated U87 cells. **(C)** Gene ontology (GO) enrichment analysis of the differentially expressed downregulated genes in U87 cells treated with sγPNA-10b/BNP in comparison to the control (untreated cells).

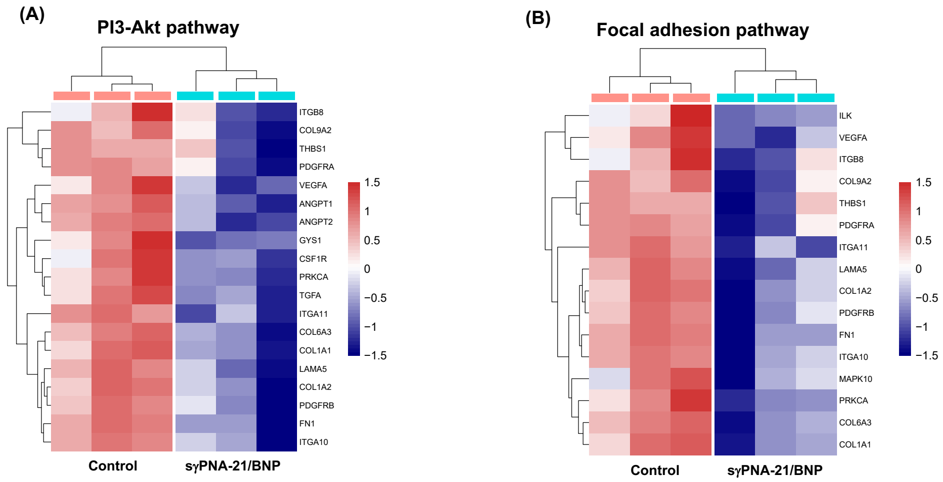


**Fig. S13. Heatmaps of PI3-Akt and focal adhesion pathway after miR-21 knockdown in U87 cells.** The heatmaps of differentially expressed downregulated genes in U87 cells treated with sγPNA-21/BNP, that are associated with **(A)** PI3-Akt pathway, and **(B)** focal adhesion pathway.

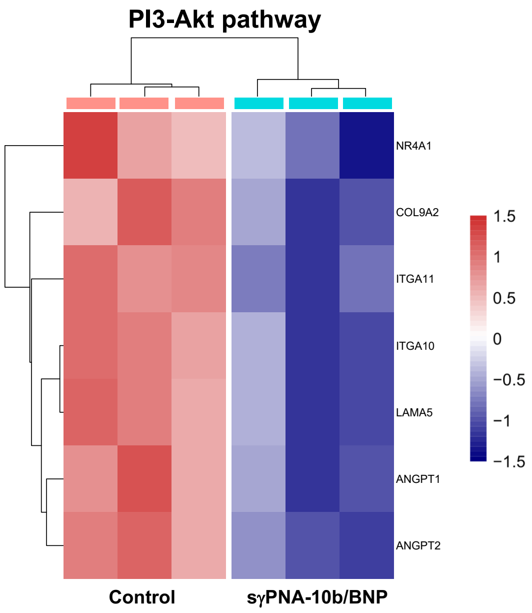


**Fig. S14. Heatmaps of PI3-Akt pathway after miR-10b knockdown in U87 cells.** The heatmaps of differentially expressed downregulated genes in U87 cells treated with sγPNA-10b/BNP, that are associated with PI3-Akt pathway.

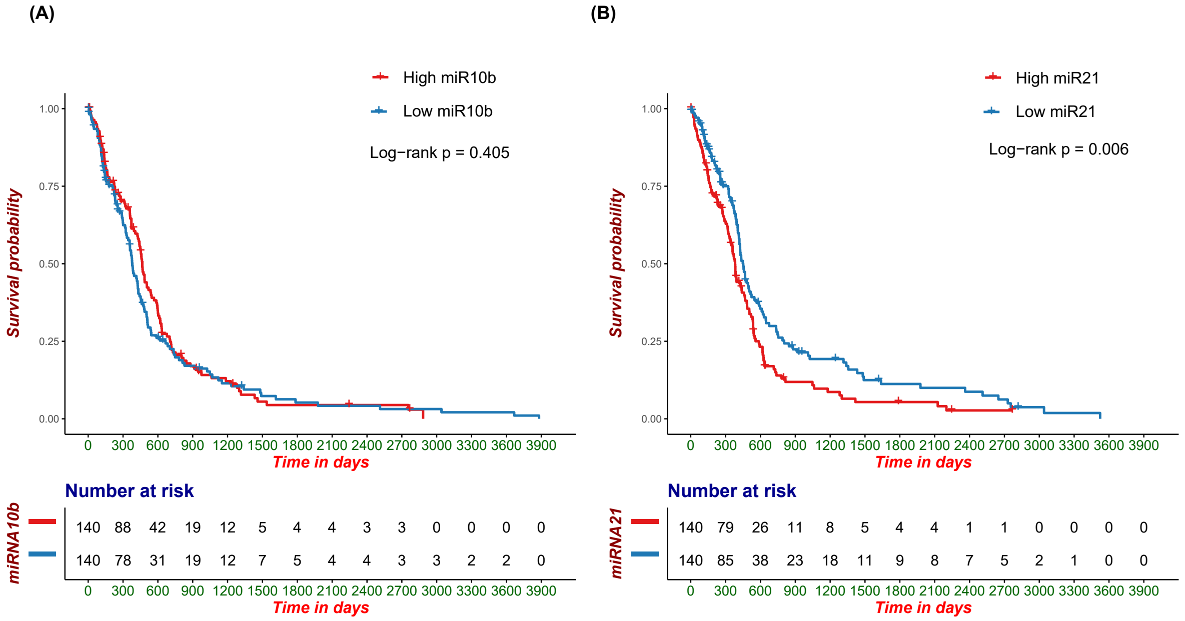


**Fig. S15. The correlation of survival probabilities and levels of miR-10b or miR-21. (A)** The survival probability of GBM patients in association with the levels of miR-10b. **(B)** The survival probability of GBM patients in association with the levels of miR-21.

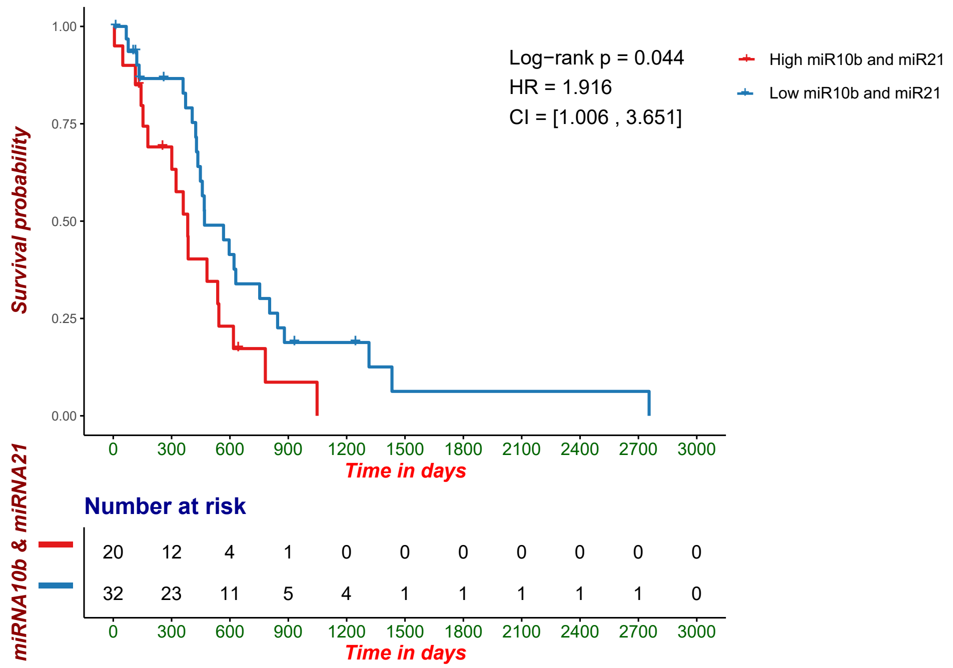


**Fig. S16. The correlation of survival probabilities and levels of both miR-10b and miR-21.**

The survival probability of GBM patients in association with the levels of miR-10b and miR-21.

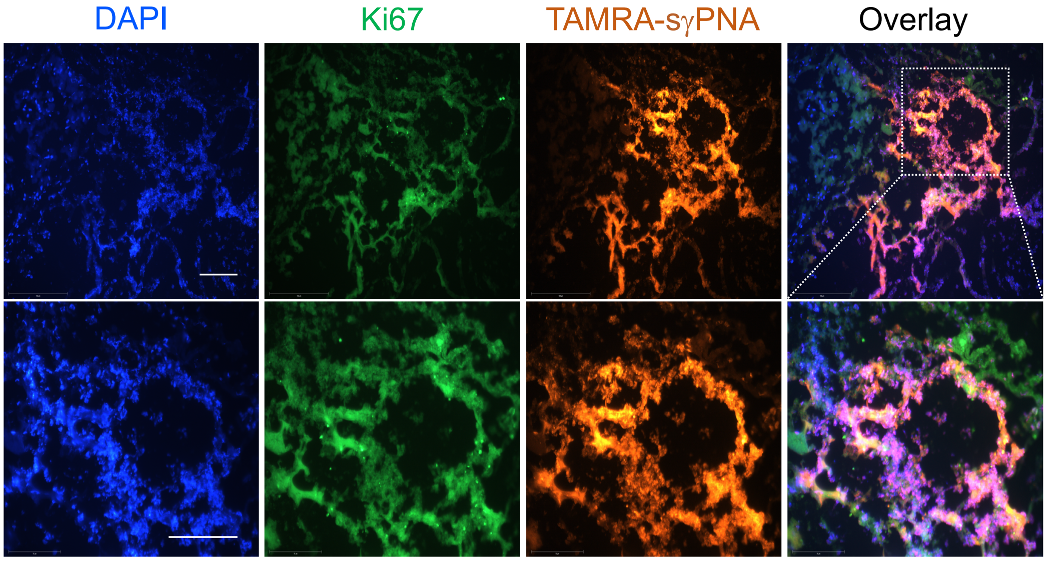


**Fig. S17. Immunostaining of ki67 in TAMRA-sγPNA /BNP treated mice (CED) after one day.** Blue indicates nucleus, green indicates Ki67, orange indicates TAMRA-sγPNA. Scale bar in top panel is 150 μm and bottom panel is 75 μm.

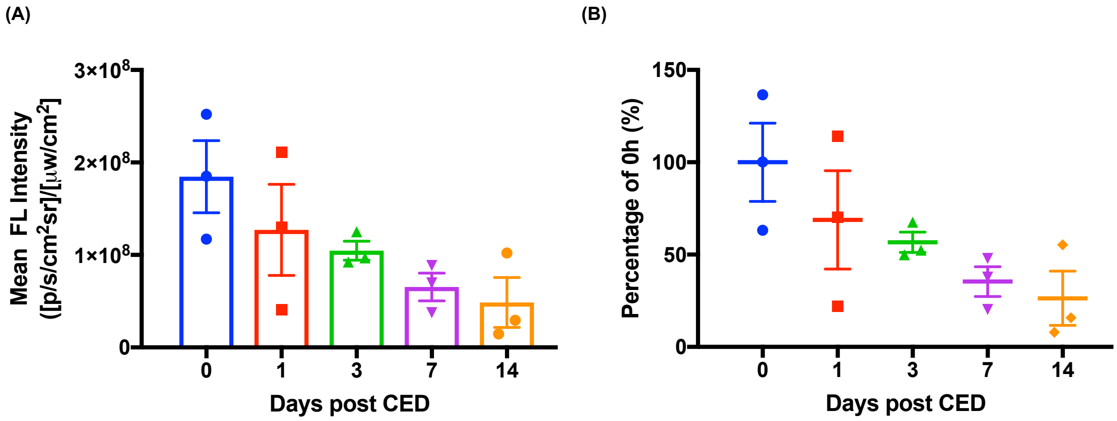


**Fig. S18. Mean fluorescence intensity of TAMRA-sγPNA in the brain. (A)** Mean fluorescence intensity of TAMRA-sγPNA in the brain at different time points and **(B)** percentage of day 0. Data are expressed as mean±sem, n=3.

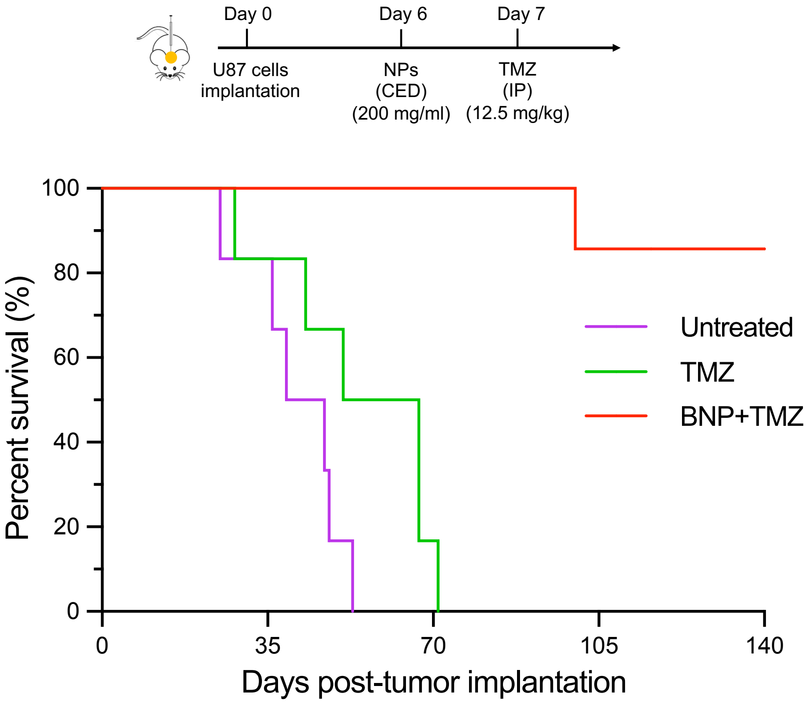


**Fig. S19. Survival study of animals treated with lower dose of TMZ (12.5 mg/kg) and combination treatment with TMZ and sγPNA/BNP**.

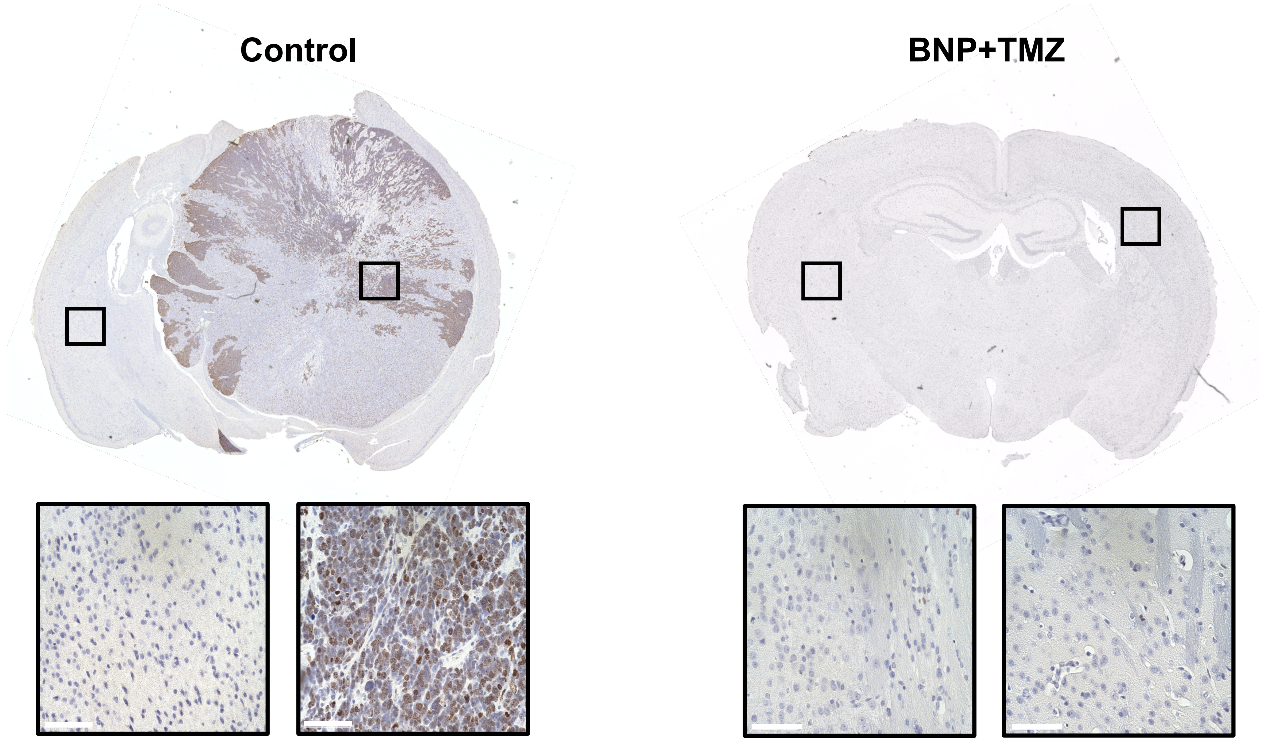


**Fig. S20. Ki67 staining of control and sγPNA/BNP+TMZ treated U87 tumor-bearing mice brain at the end of the survival study.** Scale bar, 75 µm.

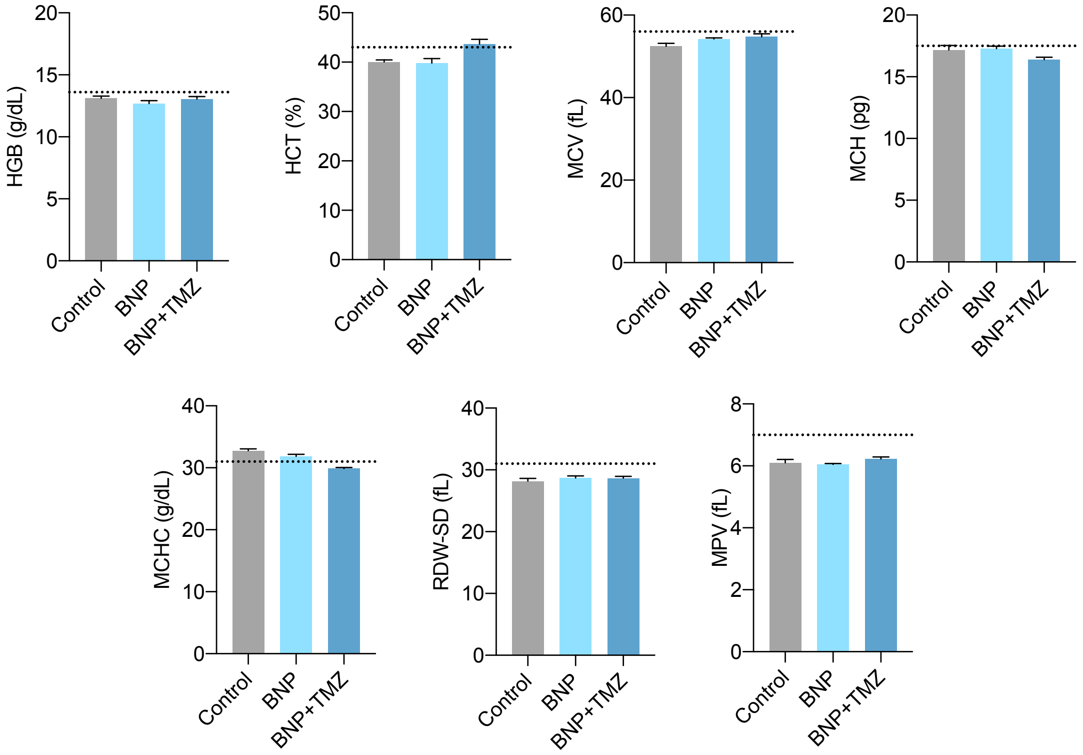


**Fig. S21. *In vivo* toxicity analysis of BNP and BNP+Temozolomide (TMZ).** The complete blood count analysis of blood samples from mice treated with BNP and BNP+TMZ after 48 hours of treatment. Results are represented as mean±sem (n=3 mice/group). HCT: Hematocrit, HGB: Hemoglobin, MCV: Mean Corpuscular Volume, MCH: Mean Corpuscular Hemoglobin, MCHC: Mean Corpuscular Hemoglobin Concentration, RDW-SD: Red Cell Distribution Width, MPV: Mean Platelets Volume.

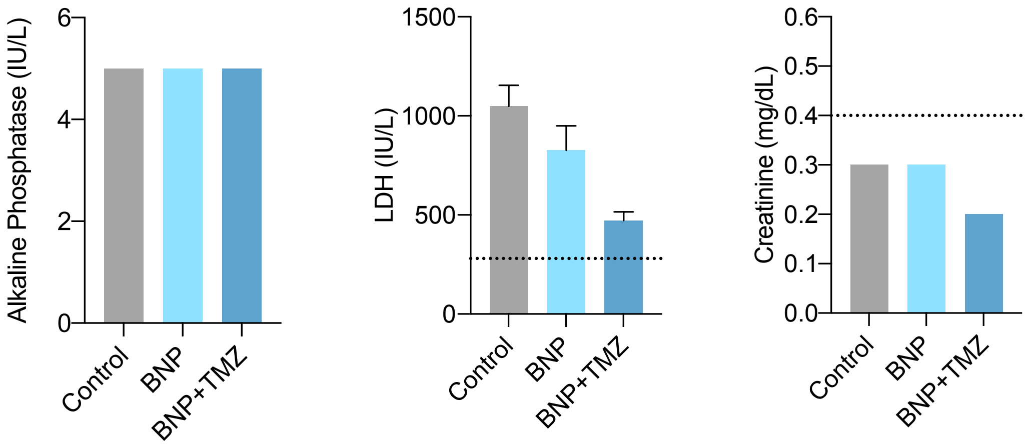


**Fig. S22. The biochemistry analysis of blood samples from mice treated with BNP and BNP+TMZ after 48 hours of treatment**. Results are represented as mean±sem (n=3 mice/group). LDH: Lactate dehydrogenase.

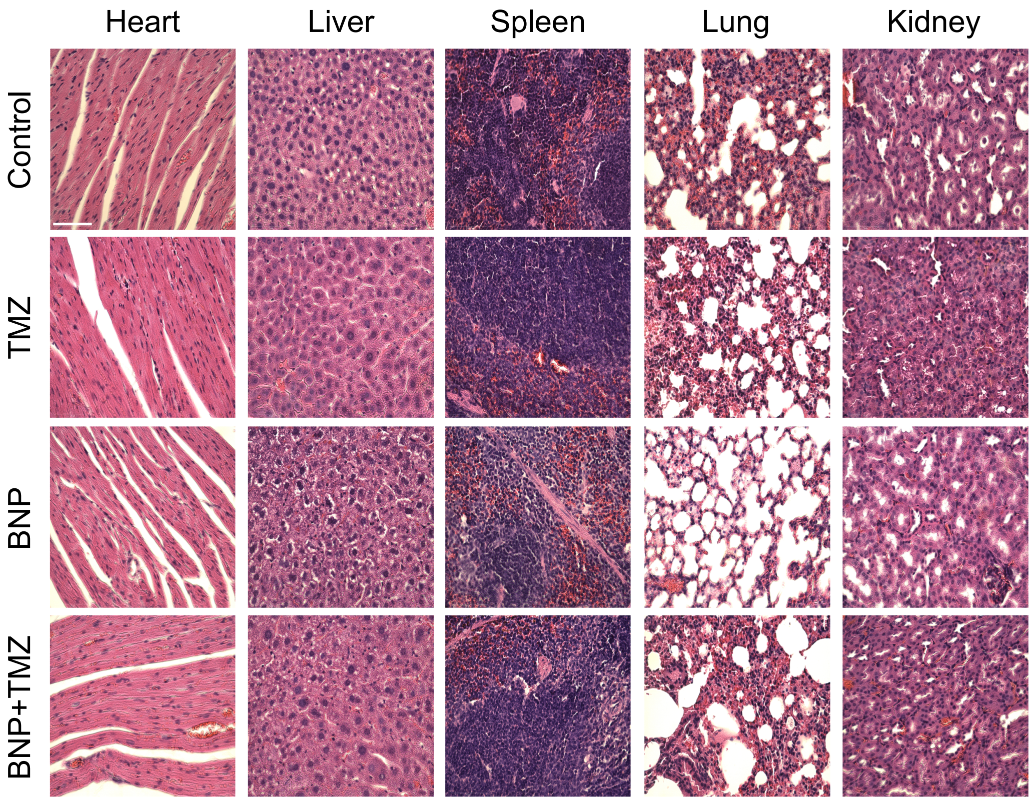


**Fig. S23. H&E staining of major organs including heart, liver, spleen, lung, liver, and kidney in all groups of mice from survival study.** The scale bar is 75 μm.

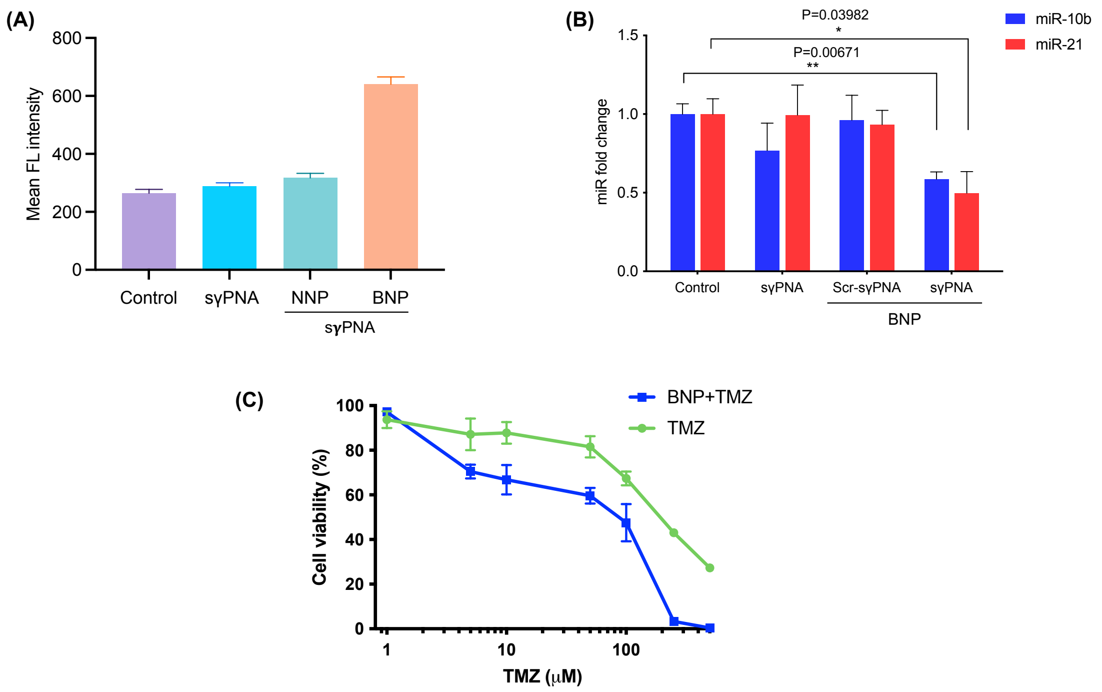


**Fig. S24. Cellular uptake and efficacy of sγPNA in G22 cells (patient derived glioblastoma cells).** (A) Quantification of cellular uptake of sγPNA in G22 cells via flow cytometry analysis. (B) The levels of miR-10b and miR-10b in G22 cells (patient derived glioblastoma cells) treated in vitro with sγPNA (sγPNA-10b + sγPNA-21) and BNPs containing sγPNAs and scr-sγPNAs. (C) Cell viability of G22 patient derived glioma cells treated by BNP and increasing doses of TMZ for 72 h. BNP are physical mixture of sγPNA-21 BNP and sγPNA-10b BNP. BNP indicates PLA-HPG-CHO nanoparticles.

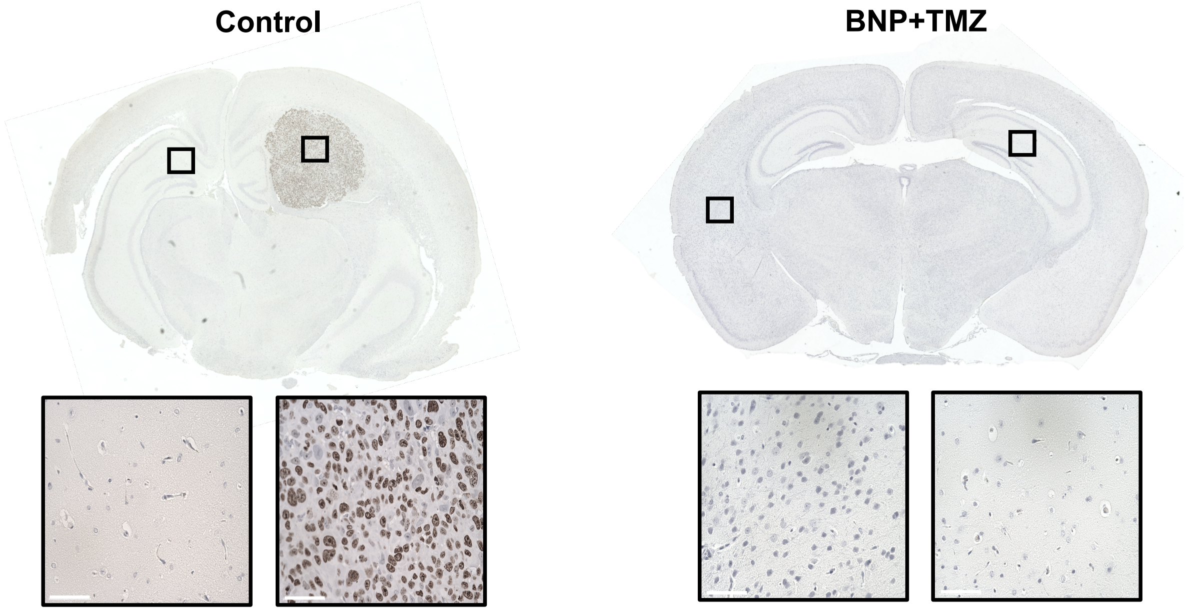


**Fig. S25. Ki67 staining of control and sγPNA/BNP+TMZ treated G22 tumor-bearing mice brain at the end of the survival study**. Scale bar, 75 μm.

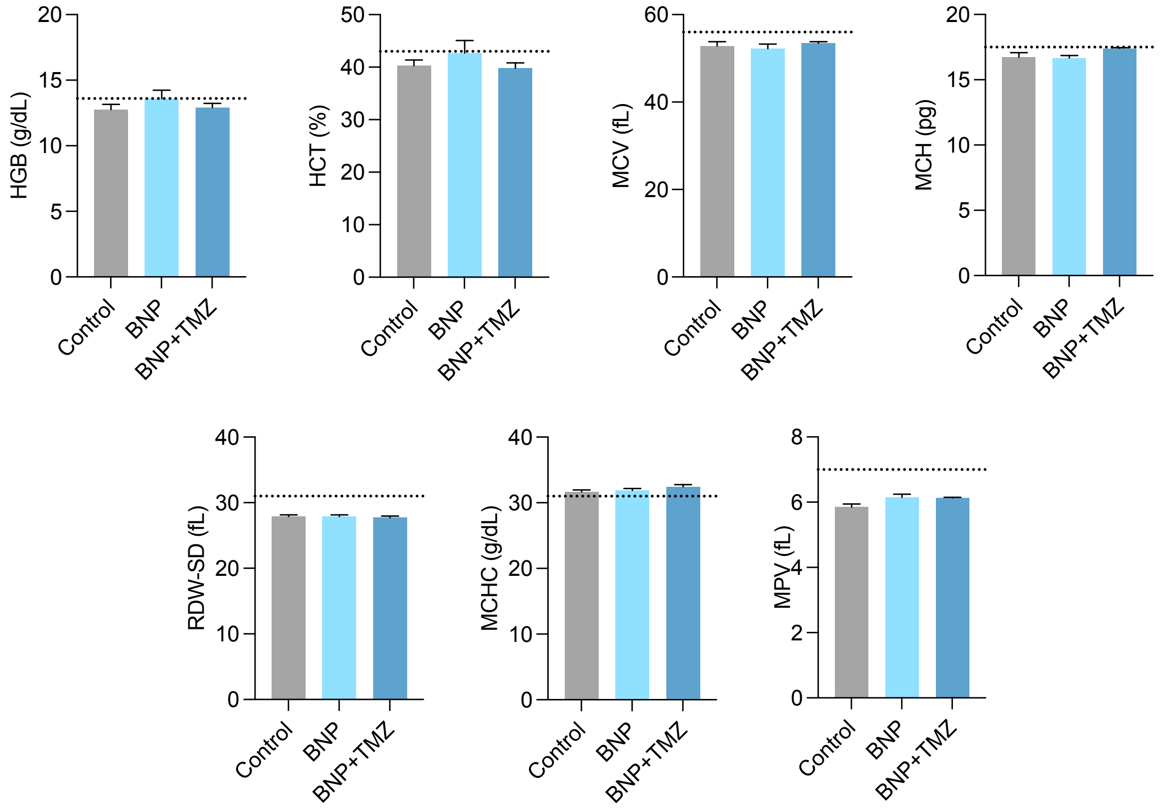


**Fig. S26. The complete blood count analysis of blood samples from G22 PDX mice model treated with BNP and BNP+TMZ after 48 hours of treatment.** Results are represented as mean±sem (n=4 mice/group). HCT: Hematocrit, HGB: Hemoglobin, MCV: Mean Corpuscular Volume, MCH: Mean Corpuscular Hemoglobin, MCHC: Mean Corpuscular Hemoglobin Concentration, RDW-SD: Red Cell Distribution Width, MPV: Mean Platelets Volume.

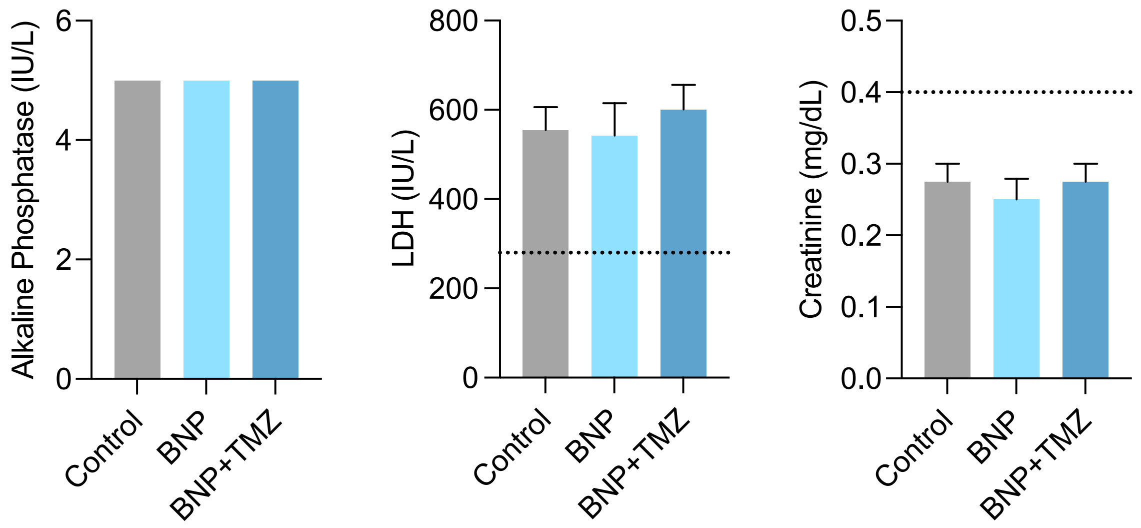


**Fig. S27. The biochemistry analysis of blood samples from G22 PDX mice model treated with BNP and BNP+TMZ after 48 hours of treatment.** Results are represented as mean±sem (n=3 mice/group). LDH: Lactate dehydrogenase.

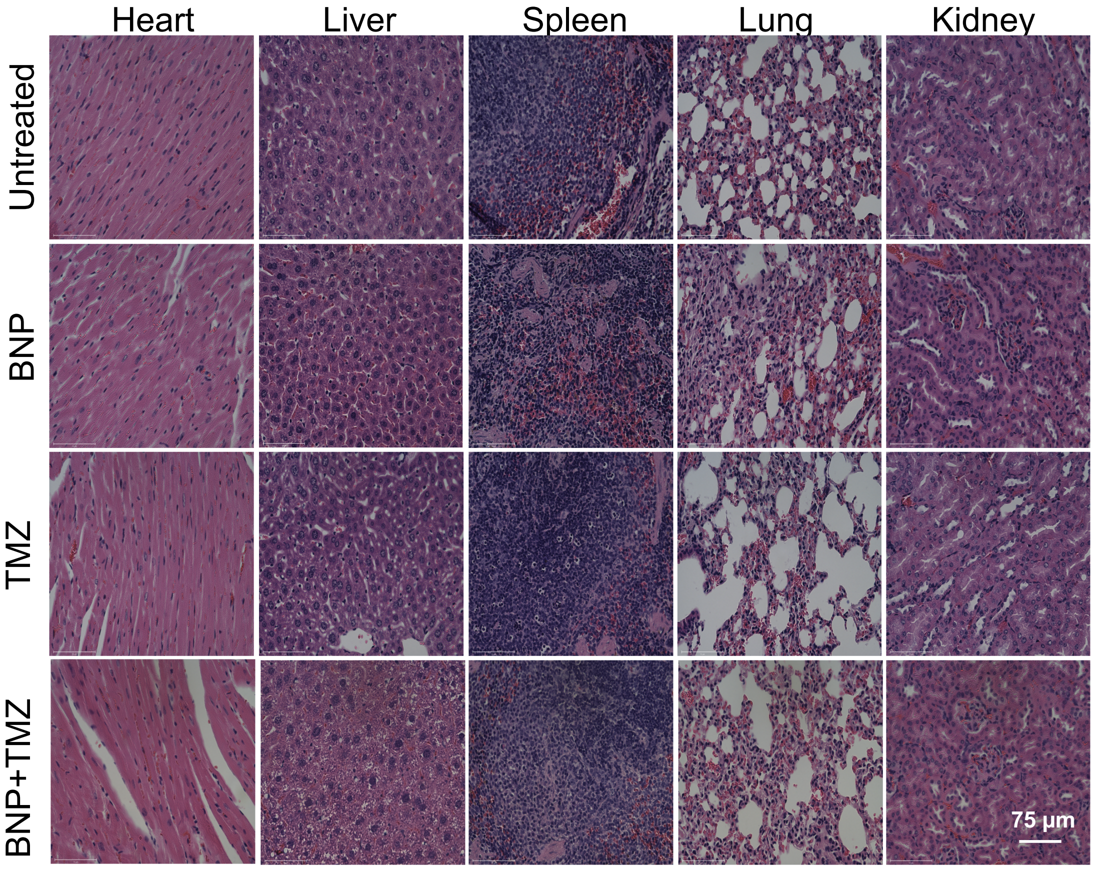


**Fig. S28. H&E staining of major organs including heart, liver, spleen, lung, liver, and kidney in all groups of mice from survival study in G22 PDX mice model of glioblastoma.** Scale bar, 75 µm.

**Table S1. Molecular weight (MW) of perfect match PNAs used in the study**. MW was determined by MALDI-TOF analysis.

| **PNA** | **Calculated MW (Daltons, Da)** | **Observed MW (Daltons, Da)** |
|---|---|---|
| sγPNA-21 | 3688 | 3693 |
| sγPNA-10b | 3704 | 3734 |
| PNA-21 | 6654 | 6680 |
| PNA-10b | 7014 | 7015 |

**Table S2**. **The synergistic effect of γPNA/BNP and TMZ combination treatment on U87 cells.** According to additive model, a ratio between the observed and the expected viability of tumor cells was calculated for the combination treatment and a ratio less than 0.8 was considered to be synergistic.

| **Treatment** | **Cell viability (% of control)** | **Cell viability/100** |
|---|---|---|
| BNP (0.5 mg/mL) | 80% | 0.8 |
| TMZ (20 μM) | 66% | 0.66 |
| BNP+TMZ combination-expected | 0.8*0.66 | 0.528 |
| BNP+TMZ combination-observed | 34% | 0.34 |
| Observed/Expected ratio | | 0.644 |
| 0.644<0.8→ Synergism | | |
